# Amira: detection of AMR genes directly from long reads using gene-space *de Bruijn* graphs

**DOI:** 10.1101/2025.05.16.654303

**Authors:** Daniel Anderson, Leandro Lima, Trieu Le, Louise Judd, Ryan Wick, Zamin Iqbal

## Abstract

Accurate detection of antimicrobial resistance (AMR) genes is essential for the surveillance, epidemiology and genotypic prediction of AMR. This is typically done by generating an assembly from the sequencing reads of a bacterial isolate and running AMR gene detection tools on the assembly. However, despite advances in long-read sequencing that have greatly improved the quality and completeness of bacterial genome assemblies, assembly tools remain prone to large-scale errors caused by repeats in the genome, leading to inaccurate detection of AMR gene content and consequent impact on resistance prediction. In this work we present Amira, a tool to detect AMR genes directly from unassembled long-read sequencing data. Amira leverages the fact that multiple consecutive genes lie within a single read to construct gene-space *de Bruijn* graphs where the *k*-mer alphabet is the set of genes in the pan-genome of the species under study. Through this approach, the reads corresponding to different copies of AMR genes can be effectively separated based on the genomic context of the AMR genes, and used to infer the nucleotide sequence of each copy. Amira achieves significant improvements in genomic copy number recall and nucleotide accuracy, demonstrated through objective simulations and comparison with alternative read and assembly-based methods on samples with manually curated truth assemblies. Applied to a dataset of 32 *Escherichia coli* samples with diverse AMR gene content, Amira achieves a mean genomic-copy-number recall of 98.4% with precision 97.9% and nucleotide accuracy 99.9%. Finally, we show that Amira consistently detects more true AMR genes across all *E. coli*, *K. pneumoniae* and *E. faecium* nanopore datasets from the ENA (n=8580, 2448 and 415 respectively) than an assembly-based approach.

## Introduction

Antimicrobial resistance (AMR) in bacteria is a significant global health challenge and a major threat to modern medicine [1]. The development and spread of AMR is primarily driven by genetic changes, typically spontaneous mutations or the acquisition of AMR genes from other bacteria. In many cases, the presence of an AMR gene or SNP is sufficient to confer resistance to a single or a broad spectrum of antimicrobials [2]. However, some evidence has demonstrated dosage-dependent effects of AMR gene copy-number variation on minimum inhibitory concentration (MIC) [3, 4]. Therefore, accurate identification of AMR gene content, including copy number, is crucial for the surveillance of AMR, and for predicting and quantifying resistance.

Accurately determining AMR gene copy number in bacteria is challenging due to the frequency of duplications and the resulting complexity in the *k*-mer based *de Bruijn* graphs or overlap graphs that are used in genome assembly. Bacteria generally contain a chromosome and a number of plasmids, and both often carry mobile genetic elements (MGEs), such as transposons or insertion sequences [5]. Many MGEs can self-replicate, excise, and integrate elsewhere in the genome while carrying cargo genes such as those associated with AMR. This can lead to large regions of sequence being shared within the same or between different DNA molecules and an amplification of certain AMR genes. Fundamentally, genome assemblers can only resolve these repetitive regions if there are enough sequencing reads that exceed the length of the repeats, and prior work has shown that on average, reads need to be at least 7kb to resolve bacterial genomes [6]. Furthermore, uneven sequencing depth across the genome and the presence of plasmids with a cellular copy number far exceeding that of the chromosome can break assumptions of genome assemblers. Therefore, fragmentation is particularly likely around complex MGE-mediated rearrangements that can occur in close proximity to multi-copy AMR genes, even with long reads. Short read Illumina data (with read lengths *<*300bp) are fundamentally incapable of resolving such regions.

Numerous tools have been developed to identify genes or alleles known to be associated with AMR [7–12] and these typically require sequence assembly as a prerequisite. Long reads generated using Oxford Nanopore Technology and PacBio sequencing platforms are often longer than the size of these repeating units, but large-scale errors can still occur in complex regions of the assembly graph, leading to duplications being collapsed or sequence being absent from the assembly entirely [13–15]. To overcome the limitations of requiring a perfect assembly, several approaches have been developed to genotype AMR genes directly from sequencing reads [8, 11, 12]. However, all of these improperly handle multi-copy AMR genes, essentially detecting the presence of single copies of allelic variants only.

We set out to design an approach which would differentiate the copies of multicopy AMR genes by leveraging the organization of the genome. Our idea was to use a *de Bruijn* graph, a workhorse of assembly, with the alphabet consisting of blocks of sequence, rather than nucleotides. There were two key requirements for how the blocks should be defined. First, the definition of a block should be robust to some sequence variation, allowing one to identify two imperfect repeats on the genome as “the same” block, and be robust to SNP and INDEL errors in noisy long reads. Second, the blocks should be sufficiently short to allow multiple blocks to be captured within a single long read (which often span tens of thousands of nucleotides in length), allowing multiple *k*-mers to be detected on a single read. Simply defining blocks as genes (and ignoring intergenic regions) will satisfy both of these constraints.

There are two major advantages to this approach. First, almost all genes in a bacterial genome are unique, which means almost all *k*-mers will be unique, and the graphs will be far simpler than nucleotide-space *de Bruijn* graphs. Second, if we encode known alleles of each gene into a pan-genome graph, reads can be mapped to mosaics of all known versions of a gene simultaneously using Pandora [16]. When reads are mapped to the pan-genome graph of the species under study, the reads which map to copies of a gene in different contexts will be naturally separated, and could then be assembled separately. How to properly handle tandem copies of genes, or more complex situations is a core aspect of this study.

In this work we present Amira, a tool to genotype AMR genes directly from long read sequencing data. As outlined above, Amira uses the genomic context of AMR genes to differentiate and cluster the reads corresponding to different copies of multi-copy genes and obtain the alleles of each copy. We show that Amira achieves improvements in AMR gene detection and allele accuracy for both single and multi-copy AMR genes compared to long read-only assembly- and read-based methods.

## Results

### Overview

We developed a method, implemented in a tool called Amira, for detecting multi-copy AMR genes directly from long read sequencing data. The first step is to identify an ordered list of the genes on long read sequences using a reference pan-genome of genes. Second, to construct a *de Bruijn* graph (DBG) in gene space and apply correction methods to the graph and the underlying reads (Fig.1a-c). Finally, to cluster the reads pertaining to different copies of the same AMR gene by their path in the graph and obtain the nucleotide sequences of each copy (Fig.1d-g).

### Gene *de Bruijn* graph construction

In our approach, Pandora is used to identify genes along a read. Pandora uses a species-specific pan-genome reference graph (panRG), which is constructed from a representative set of reference genomes, and a set of known AMR genes, using make prg and Pandora [16]. We map the reads to the panRG and store an ordered list of gene identifiers, directions and coordinates on each read, which are used to build a gene-mer *de Bruijn* graph. A read correction method is applied that removes dead ends and merges paths through bubbles in the graph that originate from the same underlying nucleotide sequence. The graph is then rebuilt from the corrected reads and we iteratively repeat the process until there are no more corrections (see Methods for further details).

### Assignment and assembly of reads to different copies of a gene

We then iterate through each unique AMR gene that has been identified by Pandora and for each, go through a 2-step clustering process, with the aim of separating the reads containing different copies of the gene. In the first step, reads are clustered based on the immediate genomic context that is 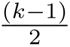 genes upstream or downstream of the AMR gene or a tandem array of AMR genes. In the second step, the full length of the reads are used to infer and cluster reads by the minimum-length genomic context that is necessary to differentiate one copy of the gene from another (see Fig. 1d; further details in Methods). Each cluster then corresponds to a genomic copy of the AMR gene. The cellular copy number (which sums the copy number, relative to the chromosome, of the molecules bearing each genomic copy of a gene) is then estimated for each genomic copy of a gene and Racon [17] is used to polish the reference allele for the gene that is most similar to the allele found in the cluster of reads. Finally, a TSV is output that summarizes the details of each copy of each AMR gene identified in the sample.

### Constructing species-specific reference pan-genomes of genes

We construct a panRG from the gene annotations of single-species reference assemblies and supplement it with 4056 plasmid-specific genes [18] (to provide additional context genes for AMR genes on plasmids) and 7062 acquired AMR gene reference alleles from the NCBI Bacterial Antimicrobial Resistance Reference Gene Database (Accession PRJNA313047) [7], 40 of which are filtered by our quality control. We then use make prg and Pandora to construct a pan-genome reference graph (panRG) from multiple sequence alignments of orthologous clusters of the annotations. The panRGs for *Escherichia coli*, *Klebsiella pneumoniae*, and *Enterococcus faecium* comprised 98796, 201978, and 163225 alleles, respectively, distributed across 11665, 14173, and 7306 clusters of orthologous genes.

**Fig. 1.**
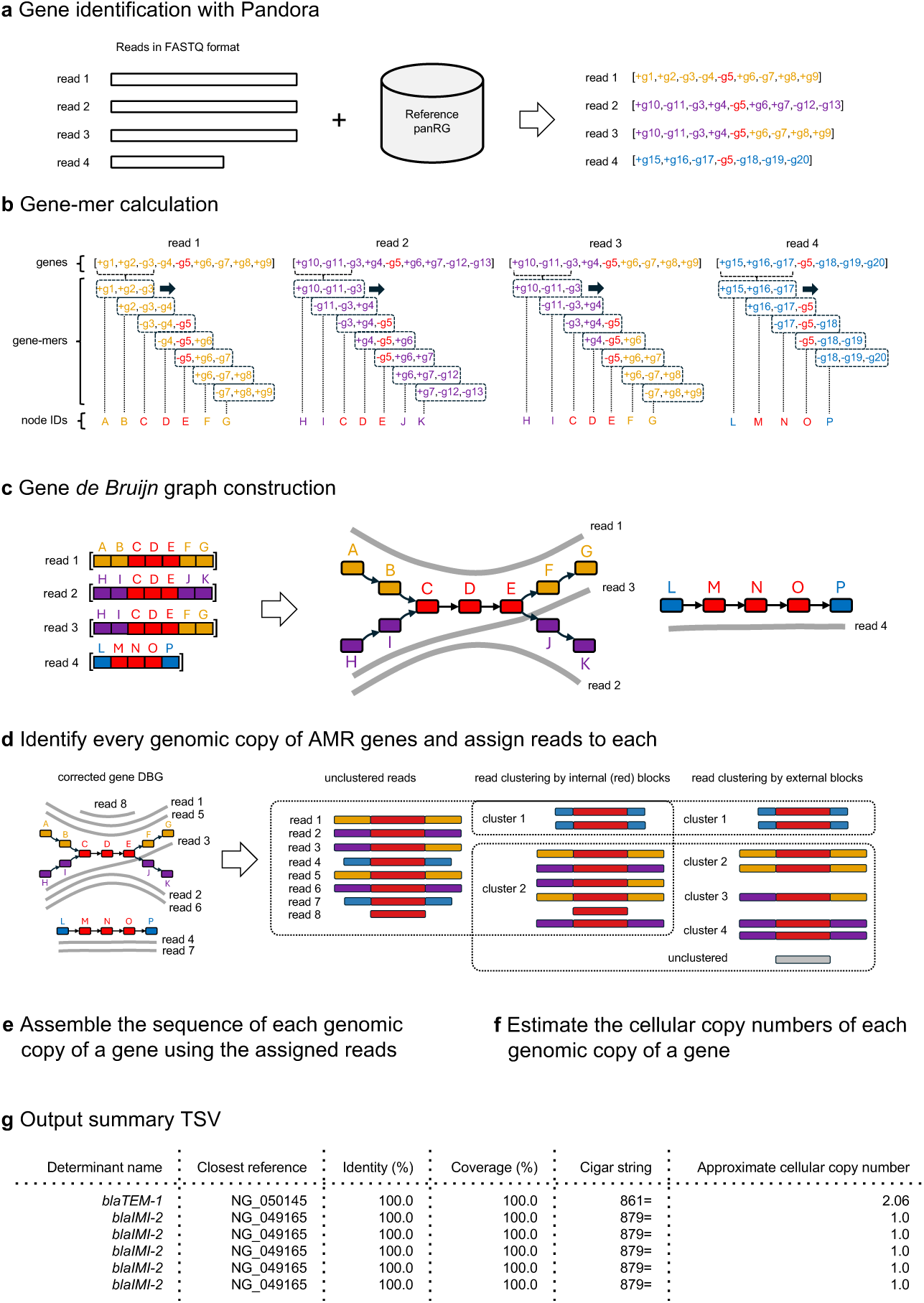
An outline of the Amira approach. (a) First, gene identifiers, orientations and coordinates are detected on long read sequences using Pandora [16]. (b & c) A DBG is constructed in gene space from the gene calls, where each node corresponds to k adjacent genes. The gene DBG is corrected by applying an iterative correction algorithm that consists of dead end trimming and bubble popping (not shown). (d) Amira uses the corrected gene DBG to cluster the reads corresponding to multicopy AMR genes based on their path through the graph, first by clustering based on the presence of internal blocks of contiguous nodes containing AMR genes, then sub-clustering using external blocks that are upstream and downstream of the internal blocks. Red nodes contain AMR genes and orange, purple and blue are used to indicate the paths of reads through this region of the graph. (e & f) After clustering the reads, the nucleotide sequence is obtained and the cellular copy number estimated for each genomic copy of each AMR gene. (g) Finally, a TSV is output that summarizes all of the AMR genes identified in the sample, with one row per genomic copy of each gene.

### Simulations confirm that Amira resolves complex duplications of AMR genes

We simulated six scenarios to evaluate Amira and to assess the impact of read depth and length on AMR gene calling accuracy. This was done by artificially inserting real AMR-gene-containing sequences of increasing complexity into a reference *E. coli* chromosome and plasmid with no AMR genes. The scenarios were: a negative control, a single isolated AMR gene on the chromosome, a two-copy gene with one copy on the chromosome and the other on the plasmid, a tandem array of six duplicated AMR genes on the chromosome, a 54.2 kb multi-gene array on the chromosome and a 36.5 kb array on the plasmid, and finally the same as the previous scenario with an additional 37.9 kb array on the chromosome [19]. We define AMR gene genomic-copy-number recall as the proportion of true genomic copies of a gene that were identified. Recall showed a strong positive correlation with increased read depth, and maximum accuracy was achieved at 40x and 80x depth due to there being a sufficient number of reads in the longer tail of the read length distribution (Fig.2). Recall was also highly correlated with read length, particularly in scenarios 3, 4 and 5, where longer reads could compensate for lower read depth. The recall of Amira was superior to AMRFinderPlus with Flye across almost all read depths and lengths.

The precise allelic identity of AMR genes correctly identified by Amira was consistently high even at low read depths, with a mean allelic identity of 99.5% at 5x, 99.7% at 10x, 99.8% at 20x and 99.9% at 40x and 80x depth.

### Evaluation on empirical data

Since simulations do not fully reflect the error profiles, read-length distributions and AMR gene complexity of real long read sequencing data, we next evaluate how Amira performs on a dataset of 32 diverse *E. coli* with Illumina and Nanopore reads available and known AMR gene content. Ten of these samples were sequenced using R10.4.1 Nanopore flow cells and 22 using R9.4.1. The evaluation samples span across the *E. coli* phylogeny and are independent from those used to construct the reference panRG (Fig.S8). The dataset contains 18 isolates with at least one multi-copy AMR gene, with one sample containing seven copies of a *blaCMY* gene, of which six are in tandem.

The mean read length across the evaluation set for the R9.4.1 samples is 14.2 kbp and Amira identified a mean of 11.9 genes per read. For the R10.4.1 samples, the mean read length is 4.7 kbp with 4.4 genes per read.

We compared the genomic copy number and type of AMR genes found by Amira (using Nanopore data) to that pre-existing tools. The main comparator tools were AMRFinderPlus run on Nanopore-only assemblies generated with Flye [21] (Flye AMRFP) and Raven [22] (Raven AMRFP), hybrid assemblies generated with Unicycler [23] (Unicycler AMRFP), and ResFinder [8]. Genomic-copy-number recall is defined as in the simulation evaluation. Genomic-copy-number precision is defined as the proportion of identified genomic copies of an AMR gene that were present in the truth assembly. Therefore, lower precision indicates a tool is over-estimating genomic copy number or calling genes that are absent from the truth assembly. These results are presented in Figure 3.

**Fig. 2.**
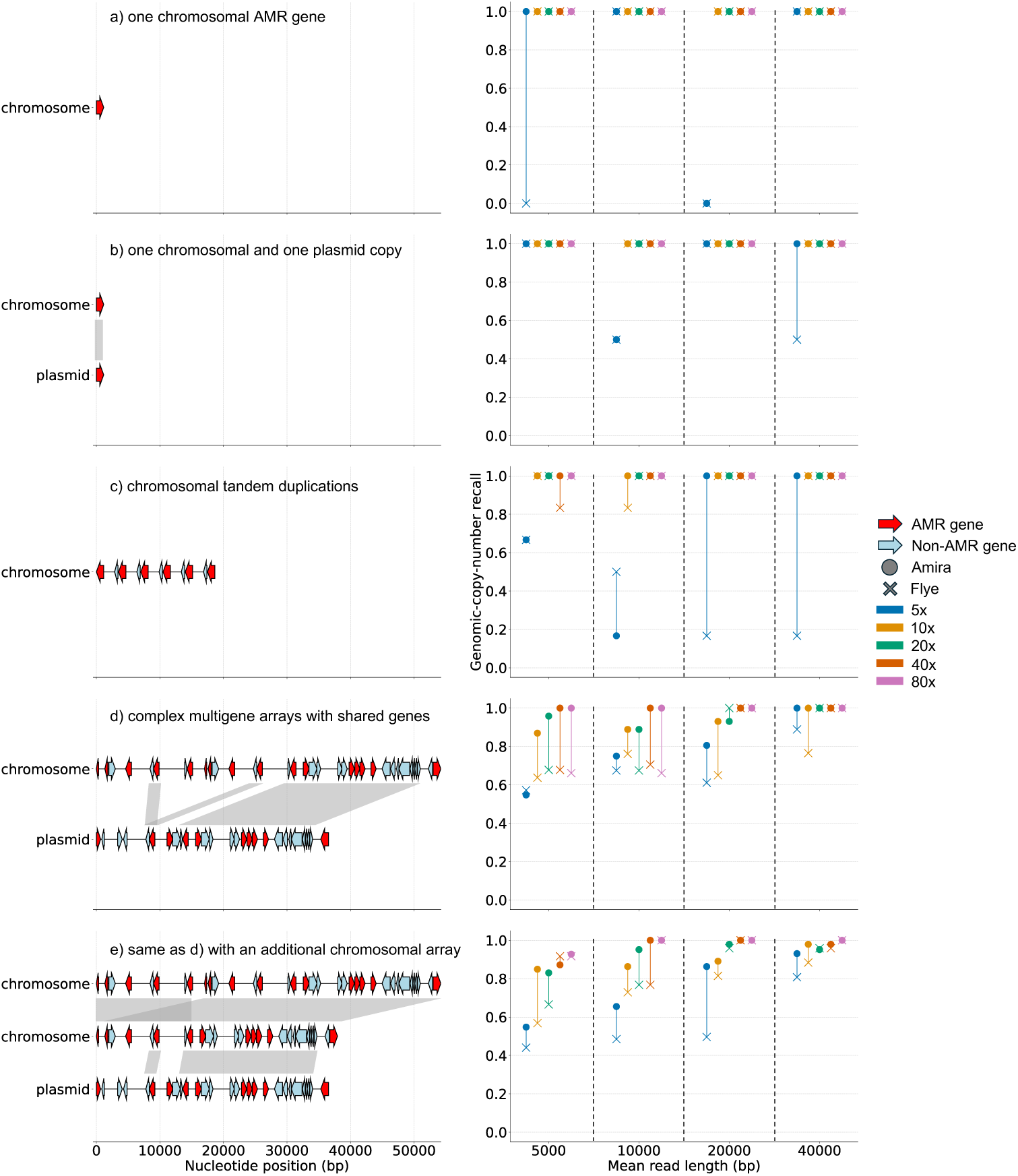
Evaluation of Amira and AMRFinderPlus with Flye in five simulated scenarios. Context plots detailing the AMR gene (red) and context gene (blue) content of five simulated scenarios (left) and the AMR gene genomic-copy-number recall of Amira and Flye with AMRFinderPlus applied to long reads simulated with Badread [20] from each scenario (right). We artificially inserted each region into the chromosome or plasmid of an artificial *E. coli* assembly containing no AMR genes, and applied Amira and Flye with AMRFinderPlus to the reads simulated from each simulated assembly. The x-axis of the left-hand plots shows the length of the inserted region in base pairs, and the y-axis shows which contig in the reference each region was inserted into. The x-axis of the right hand plots show the mean simulated read lengths (5 kb, 10 kb, 20 kb and 40 kb), the y-axis the recall for each genomic copy of an AMR gene. Each color indicates the simulated read depth (5x, 10x, 20x, 40x or 80x).

Amira achieved the highest genomic-copy-number recall on both R9.4.1 and R10.4.1 data, with 97.9% and 99.5% recall respectively, and precisions of 97.1% and 99.5% (see Fig.3a; a perfect tool would be plotted in the top right corner). Notably, the results on R10.4.1 were achieved despite a much shorter mean read length than the reads in the R9.4.1 dataset. By contrast, the performance of the other tools varied by Nanopore flow cell technology. Flye AMRFP and Raven AMRFP had low recall (71.4% and 75.6%) and high precision (100% and 98.2%) for R9.4.1, but high recall (96.0% and 93.4%) and precision (100% and 93.7%) for R10.4.1. Unicycler AMRFP had high recall (97.0%) and precision (100%) on R9.4.1 and low recall (89.2%) with high precision (98.0%) for R10.4.1. Resfinder had low recall on both datasets (71.8% and 86.4%), with higher precision on the R9.4.1 samples (99.8%) than the R10.4.1 samples (88.3%). Figure 3b shows the correlation between the cellular copy number of each AMR gene genomic copy estimated by Amira compared with the “true” cellular copy number (estimated by mapping the Nanopore reads to the truth assembly).

While we developed Amira specifically to better detect multi-copy genes, we found it excelled at detecting both single-copy and multi-copy genes in this dataset, with a mean per-gene recalls of 99.6% and 97.7% respectively (Fig.3d).

The accuracy of the nucleotide sequences for genes correctly called by Amira was consistently high for both the R9.4.1 and R10.4.1 datasets, with mean nucleotide identities of 99.9% and 99.9% respectively, compared to 98.4% and 99.6% for Flye AMRFP, 99.3% and 99.6% for Raven AMRFP, and 99.6% and 99.6% for Unicycler AMRFP (Fig.3c). The mean wall-clock time of Amira was 1,033 seconds (min 605, max 1,692) with a peak RAM of 6.1 GB (min 4.4, max 12.5) using 4 CPUs (Table.S2).

### Amira consistently detects more true AMR genes in the ENA than assembly

After finding Amira had higher recall of single- and multi-copy AMR genes than assembly-based methods, we followed up with a large-scale comparison across all available nanopore data in the ENA from three species: *Escherichia coli* (n=8580), *Klebsiella pneumoniae* (n=2448) and *Enterococcus faecium* (n=415). We compare Amira with Flye AMRFP, as used in the previous section, but here it is not possible to use manually curated truth assemblies, so we instead post-hoc confirmed whether a gene had been correctly detected (present/absent) by mapping reads to the catalogue of resistance genes.

For each gene, Amira consistently detected more samples that contained that gene than Flye AMRFP across the three species (and we confirmed these identifications were correct). On the *E. coli* dataset, Amira identified more gene-positive samples for 76.5% (101/132) of the AMR genes, and identical numbers for 13.6% (18/132) (Fig. 4a). The corresponding figures for *K. pneumoniae* were 71.2% (74/104) and 13.5% (14/104) (Fig.4b), and for *E. faecium* they were 79.7% (59/74) and 18.9% (14/74) (Fig.4c).

**Fig. 3.**
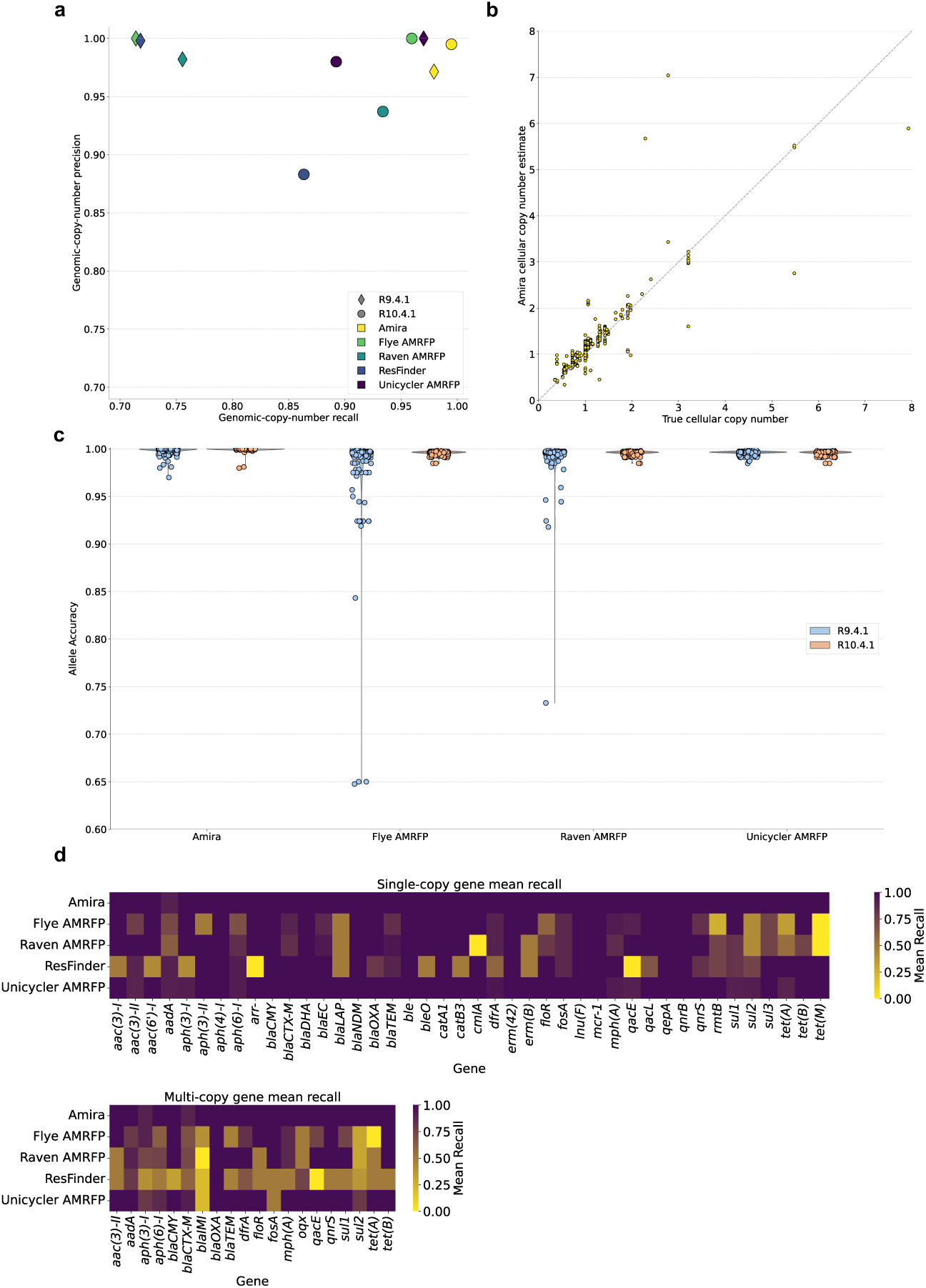
Evaluation of Amira and alternate tools for the curated assemblies of 32 *E. coli*. (a) Genomic-copy-number recall and precision on the R9.4.1 and R10.4.1 samples. Note the axes run from 0.7 to 1.0. (b) Amira cellular-copy-number estimates for each copy of each sample, compared to the true cellular copy number for the gene copy (with the truth estimated by dividing the mean Nanopore depth across the contig by the mean depth of the longest contig). (c) Gap-inclusive nucleotide similarities between the true nucleotide sequence for each each allele and that output by Amira or assembled by Nanopore-only assembly with Flye and Raven, and hybrid assembly with Unicycler. (d) Heatmaps showing the mean genomic-copy-number recall per method for each single and multi-copy AMR gene across the 32 test samples.

**Fig. 4.**
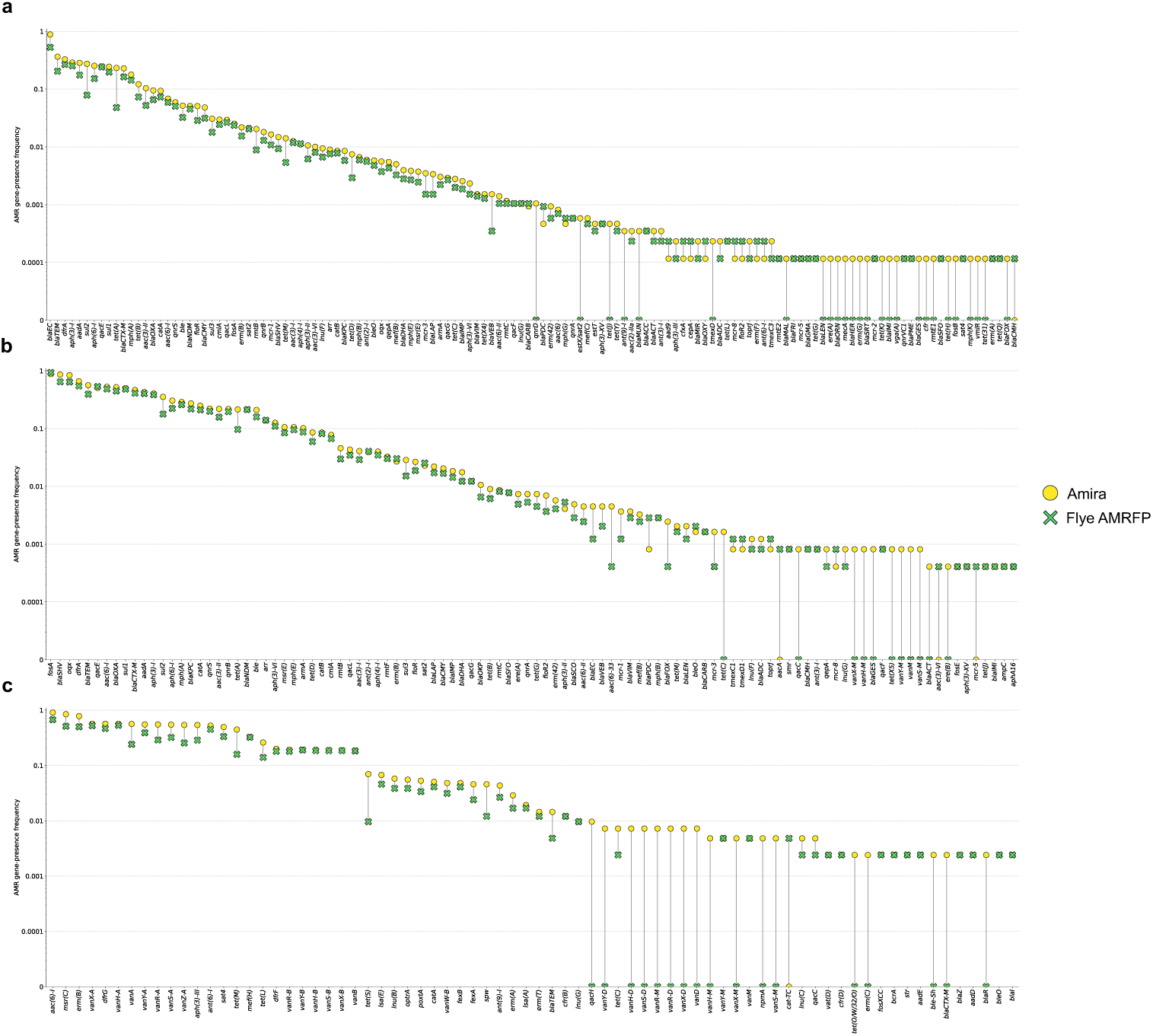
Power to detect each AMR gene in three species across the ENA: Amira vs assembly with Flye. Plots showing the proportion of samples identified as containing an AMR gene as called by Amira and AMRFinderPlus run on assemblies generated with Flye (Flye AMRFP) for (a) 8580 *E. coli*, (b) 2448 *K. pneumoniae* and (c) 415 *E. faecium* samples. Each point is a unique gene in the dataset. The y-axis indicates the proportion of samples across the dataset that contain at least one copy of the gene according to Amira or Flye AMRFP. The presence of each gene called by Amira was verified by checking the coverage of reads across the corresponding nucleotide sequence.

## Discussion

Long-read sequencing technologies have substantially improved the quality and completeness of bacterial genome assemblies. However, assemblies generated from long read sequencing data are still susceptible to large-scale errors, particularly in regions associated with duplications that are mediated by MGEs. This can result in the underestimation of AMR gene content when using assembly-based AMR gene detection tools.

We developed Amira to address this challenge. Amira leverages the full length of long read sequences to differentiate multi-copy genes by their local genomic context. Through evaluation on simulated and curated truth datasets, we have demonstrated that Amira is more sensitive in detecting the presence and the number of copies of AMR genes than competing methods. This improvement is largely driven by use of the gene as the fundamental unit. By using Pandora [16] to identify the genes on each sequencing read, Amira is robust to nucleotide-level SNP and INDEL errors, which are common in long-read sequences, as evidenced by the improvements in accuracy when applying Amira to datasets sequenced with error-prone R9.4.1 Nanopore reads. This allows the construction of significantly cleaner graphs that more accurately capture the underlying genomic structure than conventional *k*-mer based methods, leading to better resolution of gene order and assignment of reads to the correct copy of each gene. We find this gives improved nucleotide-level accuracy compared to existing assembly- and read-based methods.

Counter to our expectations, the improvement in genomic-copy-number recall, compared to assembly-based methods, is not solely due to multi-copy genes. Our interpretation is that this difference highlights the extent to which AMR gene detection from assembly is affected by contig breaks. AMRFinderPlus is able to report partially present genes, but users must manually distinguish genes that are genuinely fragmented (e.g. by MGE insertion) from artefactual fragments caused by imperfect assembly. We estimated the impact of this difference on broad-scale surveillance of AMR gene prevalence by applying Amira to the Nanopore reads of 8580 *E. coli*, 2448 *K. pneumoniae* and 415 *E. faecium* and report the proportion of samples containing at least one copy of each gene. This confirms there is consistent under-detection of AMR gene content when using assembly-based approaches (Fig. 4). However, the recall of Amira is also sometimes imperfect, particularly for multi-copy genes that share subpaths in complex regions of the graph. Although this was not an issue in the empirical evaluation of 32 samples, we expect that this resulted in the reduced gene-presence frequency that was observed for some genes in the ENA evaluation.

Simulations of progressively more challenging AMR gene arrays allowed evaluation of the relative effects of read length and depth and the intrinsic repeat structure of the genome. The genomic-copy-number recall of Amira and Flye improved with longer reads and greater sequencing depth, but we found that Amira was better able to leverage lower depth and shorter reads even in the most complex simulation scenarios. Reference bias is a well documented challenge in bacterial variant-calling methods that rely on reference sequences, which arises when a single reference incompletely represents the diversity of a sample under analysis [16, 24, 25]. Hard bias refers to the absence of entire genomic regions from a reference, often a consequence of horizontal gene transfer. With Amira, we mitigate this by detecting genes from a pan-genome reference graph (panRG) constructed from a phylogenetically representative set of genomes of the species under analysis (and currently support *E. coli*, *K. pneumoniae*, *E. faecium*, *Streptococcus pneumoniae* and *Staphylococcus aureus*). In principle, if a sample has a multi-copy AMR gene that is flanked by rare genes not present in the panRG, the flanking genes will be invisible to Amira and it may not be possible to distinguish the reads corresponding to different copies of that AMR gene. By definition, this should be a rare occurrence. However, a more significant challenge is that the plasmid pan-genome in Enterobacteriaceae is extensive and there are over six million annotations in PLSDB [26]. Rather than inflate our panRG to include huge numbers of rare plasmid genes, we chose to incorporate plasmid-specific genes from MOB-suite [18] into the panRG to expand the number of context genes that can be detected. Although this is sufficient in our dataset, this is a limitation of our approach and a future improvement could involve expanding the panRG to include genes from plasmids with high abundance or of clinical concern.

It is worth comparing our approach with that taken by wtdbg2 [27]. Wtdbg2 assembles long-read sequences by binning reads into 256 bp segments, defines *K*-bins (a sequence of *k* consecutive bins on a read), aligns the sequences of *K*-bins to merge equivalent *K*-bins, then constructs a “fuzzy-Bruijn” graph where *K*-bins are nodes. As with our approach, wtdbg2 seeks to use a block-alphabet to simplify assembly. It has the advantage that it avoids reference-bias and can be run on any sample without a predefined pan-genome; however, it also has the disadvantages that the blocks are arbitrary, and that the block size may bear little relation to the repeats in the genome. In contrast, genes are broadly conserved in bacteria, and generally provide a reliable set of blocks to define genome organisation. Further differences lie in the details of the assembly methods: wtdbg2 uses standard approaches, simplifying the graph before outputting a consensus sequence, whereas Amira infers the contexts of AMR genes by identifying paths through the graph, and leverages these contexts to cluster the reads. Recent work has found correlations between AMR gene dosage and MIC for specific gene-antimicrobial pairs [3, 4]. While MIC prediction was beyond the scope of this study, we anticipate that the more comprehensive identification of AMR gene content by Amira could enhance MIC prediction accuracy. Additionally, while Amira is designed to detect AMR genes, it also allows users to detect known and novel variants in the promoter sequences of *ampC* and *blaTEM* genes which can contribute to MIC in *E. coli* [28].

Lastly, our approach is not just limited to AMR genes and could be directly applied to estimate the genomic and cellular copy numbers of any gene of interest through minor modifications. We also introduce the gene *de Bruijn* graph that was initially proposed in [16], a data structure that moves away from conventional *k*-mers and is more robust to structural rearrangements that are frequent in bacteria. The gene DBG has the potential to be beneficial for a range of bioinformatic applications in place of *k*-mers, particularly in bacterial genome assembly, and the approach can capture structural heterogeneity even at the level of single bacterial isolates.

## Conclusion

Long read sequencing has significantly improved the completeness of bacterial genome assembly. However, long-read genome assemblies still contain errors that prevent the comprehensive identification of AMR genes. To address this, we developed Amira, a tool that effectively clusters and assembles the reads corresponding to different copies of multi-copy AMR genes using their genomic context, through the use of gene-space *de Bruijn* graphs. This new perspective, made possible by modern long read technologies, allows us to work at the level of gene organisation and provides considerable benefits. We provide pre-built pan-genome indexes for five species, and believe Amira represents a significant improvement in accuracy and accessibility of AMR gene detection.

## Methods

### Constructing species-specific panRGs

Amira relies on Pandora [16] to obtain an ordered list of gene identifiers, orientations and coordinates for each sequencing read using species-specific reference pan-genomes (abbreviated to panRGs). We developed a snakemake [29] workflow to automate the construction of these panRGs from the annotations of a set of reference assemblies. This is supplemented with the annotations of 7062 acquired AMR gene reference alleles from the NCBI Bacterial Antimicrobial Resistance Reference Gene Database (Accession PRJNA313047), and 4056 plasmid-specific alleles used by MOB-suite [18] to improve the representation of plasmid genes. The workflow begins by annotating the genes in the reference assemblies using bakta v1.7.0 [30], supplements the annotations with the AMR and plasmid genes, then clusters the annotations into orthologs using Panaroo v1.5.0 [25] `with --clean-mode sensitive --refind-mode off --remove-invalid-genes --length outlier support proportion 0.1 --merge paralogs --threshold 0.8 --len dif percent 0.0 -a pan`. All alleles less than 250 bp in length and all transposase or IS-related genes are then filtered, as these a significant source of inconsistent gene calls. After aligning the filtered clusters, we run make prg v0.5.0 with default parameters, and construct a panRG using Pandora [16] index v0.12.0, specifying `-k 15 -w 5`.

For the reference assemblies for *E. coli*, we re-assembled the Illumina and Nanopore reads used in the 20-way analysis in [16] with Hybracter v0.7.3 [31] using default parameters. For *K. pneumoniae*, we retrieved 565 hybrid assemblies and short reads from [32]. Short reads were mapped to the *K. pneumoniae MGH 78578* reference genome (Accession *CP000647.1*) with BWA v0.7.18 [33]. Variants were called using BCFtools v1.9 mpileup [34], and a phylogenetic tree was built with FastTree v2.1.11 [35]. Isolates were clustered using FastBAPS v1.0.8 [36]. 40 isolates from across the phylogeny were selected to construct the panRG (Fig.S1). For *E. faecium*, we used the 62 hybrid assemblies generated in [37] (Accession PRJEB28495).

### Constructing a read-coherent *de Bruijn* graph in gene space

Amira takes a FASTQ file of long reads as input, then uses Pandora to create a pseudo-SAM file that identifies the genes on each sequencing read. A hash table is created where the keys are read identifiers and values are an ordered list of gene identifiers, where the prefix of each gene identifier is the orientation of each gene relative to the orientation of the sequences of the gene in the panRG. A *de Bruijn* graph is then constructed in gene space (referred to as a gene DBG) using the gene calls for each read. For each read of length *n* that contains at least *k* genes, a sliding window of size *k* is moved across the list of genes in single-gene increments to generate *n → k* +1 sub-lists, referred to as “gene-mers,” that are analogous to *k*-mers used in sequence assembly. The canonical representative for each gene-mer in a read is hashed and added as a node in a graph, where the canonical is defined as the lexicographically smaller of the gene-mer and its reverse complement. An edge is then added between each pair of adjacent gene-mers and it is annotated with the orientation of the target gene-mer relative to the canonical of the source gene-mer.

### Correcting false paths in the graph

There are two types of error we expect when mapping reads to the panRG with Pandora: complete failure to detect a gene or false positive detection. Gene detection can fail for two reasons. First, if a gene present in the reads of the sample under analysis is not present in the panRG, it cannot be detected. This may occur because none of the reference genomes used to construct the panRG contain rare genes found in the sample. These genes will be consistently absent across all reads, and contribute consistent paths to the gene DBG. Second, if sequencing errors ablate too many minimizers from a read, there may too few minimizers matching to the panRG to trigger gene detection. These errors are random and contribute erroneous paths to the graph that are low coverage, so can be removed. However, there are fewer potential minimizers to detect in short genes, so the impact of sequencing errors is larger. We address these up front, by setting a minimum length threshold on genes in our panRG of 250bp, and at run-time by applying low coverage cleaning thresholds to the graph. The second major error type is false positive detection of a gene. We found these occurred systematically at loci corresponding to fragmented partial genes, particularly transposases. The frequency of such fragments is presumably due to preference of MGEs to insert near other MGEs. We avoid the problem by removing transposases from our panRG. Amira uses several methods to account for the remaining errors, first applying correction to the graph itself, then using the corrected graph to correct the underlying list of genes of each sequencing read.

Following initial graph construction, all nodes that are not contained in any reads with an AMR gene are pruned. All reads where *>*80% of the nodes along the path followed by the read have a depth equal to one are then discarded to remove reads that poorly support the graph due to error or contamination. The gene DBG is reconstructed using the remaining reads, and the dominant errors we now expect are randomly distributed and caused by Pandora failing to identify genes. If this occurs *↑ k* genes at the end of a read, the result will be a dead-end path of length *< k* and, otherwise, a low coverage alternative path of length *< k* nodes will be created. Independent errors may occur in close proximity in the graph (for example in a single read), resulting in complex regions of overlapping erroneous paths. Therefore, to obtain the cleaned gene DBG, an iterative correction algorithm is applied that consists of two stages: dead-end removal and path replacement. The dead end removal stage finds and removes all paths of length *< k* nodes, which start at a dead end, are immediately followed by a junction, and have no junctions in between. Path replacement identifies potential erroneous paths by enumerating the paths between all pairs of nodes with *>* 2 neighbors, selects the path with the highest mean node coverage as a true path, then compares this path, at the nucleotide level, to all lower coverage paths with the same terminal nodes to correct paths that in fact correspond to the same underlying nucleotide sequence. The next paragraph details this method.

Path replacement begins by defining a set of candidate nodes that have an in or out degree greater than one. A depth-first-search is then conducted between all pairs of non-identical nodes, to obtain a set of all of the paths with length *↑* 4*k*. A reduced set of paths is defined by filtering out all paths that are wholly contained within another path. The filtered paths are sorted by length in ascending order, and the paths clustered such that all paths in a cluster share a start and stop node. The clusters are processed in ascending order of maximum path length within the cluster and for each cluster, the paths are sorted by descending mean node coverage. For each path *P_i_*with index *i* in a cluster, we consider each lower coverage path *P_j_* with index *j*, where *j > i*, to be a potential false path. We extract the sets of nucleotide minimizers *M_i_* and *M_j_*from the reads for *P_i_* and *P_j_*, respectively, using sourmash v4.8.4 [38] with *k*-mer size of 11 and scaled parameter of 10. Reads covering *P_j_* are corrected to follow 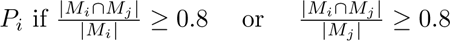.

### Assigning reads to paths

For each copy of each AMR gene in a bacterial sample, there exists a minimum length path through the corrected gene DBG that incorporates a sufficient number of context genes to make it distinct. When the genomic context immediately flanking each copy of a multi-copy AMR gene differs, only one context gene is needed. However, in cases where the number of genes within a duplicated genomic region (that contains an AMR gene) significantly exceeds the value of *k*, it is necessary to leverage the full read length to differentiate the paths.

We first outline the approach, and then describe how this is achieved in terms of algorithm and data structures. For each unique AMR gene *g* identified in the reads by Pandora, we first separate copies that occur within different immediate contexts by analyzing the reads containing *g*. For each read (treated as an ordered list of genemer nodes), we start at the first left-hand node containing *g*, and extend right until we hit the final node on the read which does not contain *g*; we term this an “internal block” and the first and last nodes of the block are called “anchors”. Reads are first clustered by the presence of internal blocks containing *g*, and we see in figure 5 that this leads to natural separation of reads 1,2,3,4 (containing internal block CDE) from read 5 (containing block MNO), despite all reads containing g5. These clusters are then sub-clustered by the paths on either side of the internal block (referred to as external blocks). In figure 5a we see that the external blocks allow us to distinguish the paths followed by reads 1 and 2 (yellow-yellow) from reads 3 and 4 (purple, purple), and from read 5 (purple, yellow). These clusters enumerate the different genomic copies of *g* present in the sample, and since Amira assigns reads to each path, the nucleotide sequences that are specific to each genomic copy can be used to assemble it.

The example in figure 5 is trivial. In reality, we expect a number of challenges with this approach. Firstly, reads may start and end at any point along a path corresponding to a unique genomic copy of a gene, so internal blocks that are partial must be filtered out (Fig.S10a). Secondly, internal blocks may be nested and a shorter block from one read may be entirely contained within a longer block from another read, but correspond to a distinct full length path (Fig.S10b). This means the anchor nodes for one read may occur within the internal block of another. Finally, a single long read may cover multiple, distant contiguous blocks of nodes containing *g*, and it is preferable to process each as an independent internal block to maximize the number of reads assigned to each (Fig.S10c).

To facilitate this, we begin by constructing a generalized suffix tree, where each ”letter” represents a node identifier, and each ”word” corresponds to the ordered list of node identifiers for the path followed by a read through the gene DBG. Each read contributes two words to the suffix tree: one for its forward orientation and the other for its reverse (specifically reverse, not reverse complement). This primary suffix tree is referred to as *T_p_*.

Internal blocks are defined for each unique AMR gene *g* in the first round of clustering as follows. Firstly, the reads containing *g* are identified and the set of anchor nodes *A* is identified, as described above, and shown in Fig 5b. A node is defined as an anchor node if it contains *g*, and appears adjacent (in any read) to a node that does not contain *g*. Every anchor node *a_i_* in *A* is queried in *T_p_*to obtain all the sublists of nodes downstream of *a_i_*, and a secondary suffix tree *T_s_* is constructed from the reverse of the suffixes (Fig.5c). Every other anchor node *a_j_* is then queried in *T_s_* to collect and cluster reads by the longest internal block they contain.

After identifying the set of internal blocks *B* for *g*, the paths of nodes external to each internal block *b_i_* in *B* are used to differentiate long duplicated regions with paths that are followed by the reads and diverge outside of *b_i_*. This is achieved by querying *T_p_* for *b_i_* and tracking the suffixes downstream of *b_i_*. Simultaneously, the reverse of *b_i_*is queried in *T_p_* and the reverse of the suffixes obtained is stored as upstream external blocks. This creates two sets for *b_i_*: one set *U_i_* corresponding to the external blocks upstream of *b_i_*, and one set *D_i_* corresponding to the downstream external blocks (Fig.5c). The following clustering approach is applied to *U_i_* and *D_i_* that infers the number and full length of the unique underlying paths supported by each external block. The elements of *U_i_* and *D_i_* are sorted by descending path length and blocks are clustered together if they are fully contained within a longer clustered block, skipping uninformative blocks that are sub-blocks of the longest block in more than one cluster (Fig.5d). Subsequently, the shortest external block in each cluster is selected as the representative for the underlying path for that cluster, and all representative blocks are stored in a set 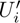 or a set 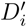 where 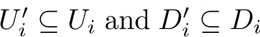. For each *b_i_*, a minimum-length node-space path is inferred by joining any cluster representative of 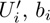, and cluster representative of 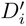 if they all share at least one read, and paths are filtered out if they are sub-paths of any longer path. The number of context genes necessary to differentiate the paths is minimized by first converting each node-space path into gene-space (*P_i_*). All possible sub-paths of *P_i_*of any length are enumerated, and any sub-path filtered out if the total number of reads it occurs in is fewer than the product of 1*/*20 and the mean node coverage, is a sub-path of any other full-length path *P_j_*, or does not contain the same number of copies of *g* as *P_i_*. The sub-path that satisfies this criteria and is covered by the most reads is selected as the final path (Fig.5e). Finally, all reads that contain each final path are collected by querying *T_p_* and two FASTQ files are generated: one file stores the full sequences of all reads containing the path, enabling estimation of the cellular copy number of the path; the other file stores the nucleotide sequences corresponding to each AMR gene copy in the path.

**Fig. 5.**
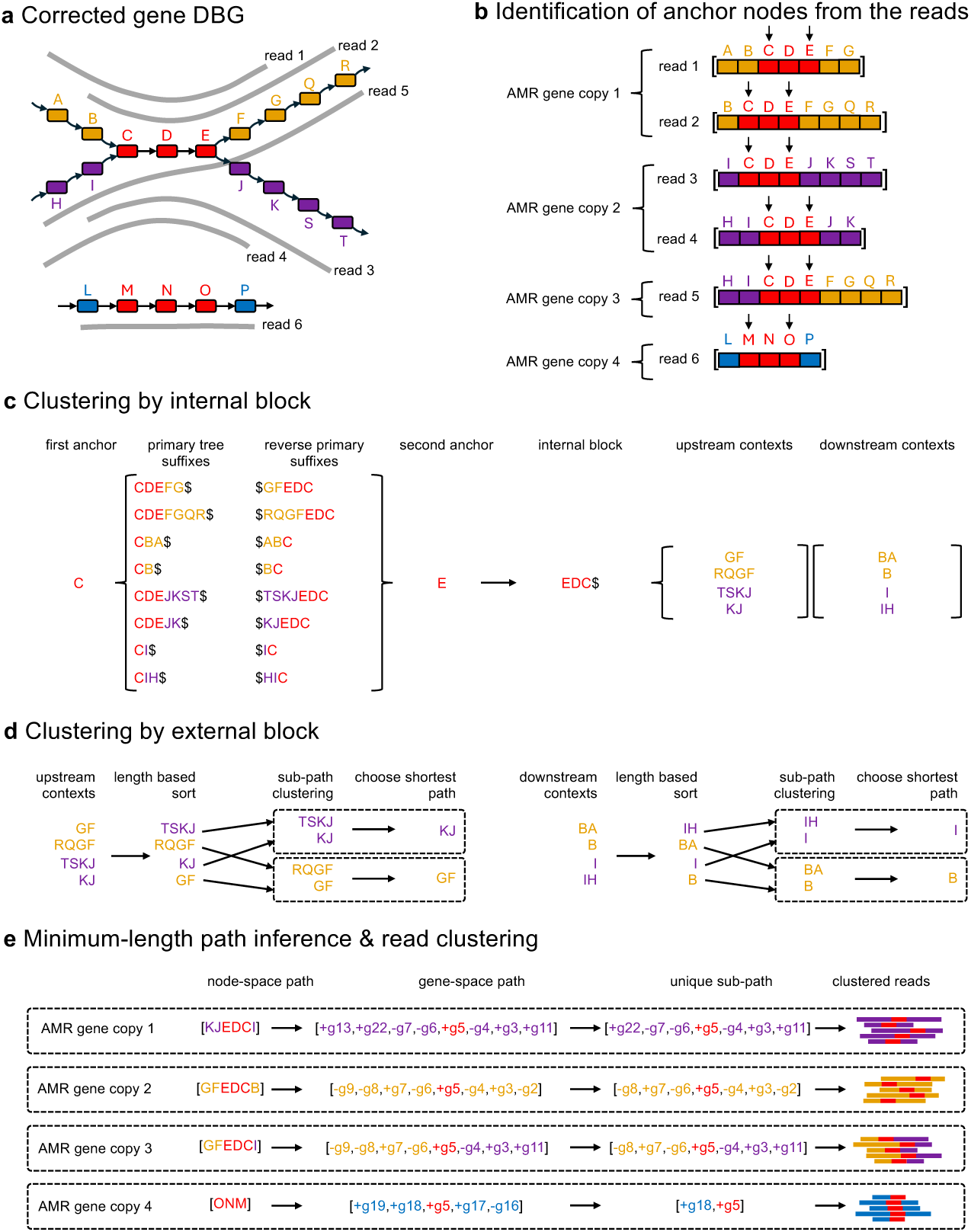
Distinguishing different copies of an AMR gene and assigning reads to each. (a) Amira compares the paths through the corrected DBG traced by the reads containing the AMR gene of interest, and uses them to infer the different copies of the gene, and their contexts. (b) The path of nodes followed by each read through the gene DBG is used to define set of “anchor” nodes for each unique AMR gene. (c) Internal blocks are defined between pairs of anchor nodes. This process begins by identifying the suffixes of the first anchor by querying a suffix tree built from the ordered list of node identifiers in the path through the gene DBG that is followed by each read. Next, the second anchor node is queried in a second suffix tree that is constructed from the reverse of the suffixes output by the initial search to obtain the internal blocks. The blocks of nodes external to each internal block are then obtained. (d) External blocks are clustered into underlying paths. External blocks are sorted from longest to shortest, and clustered with a longer block if they are entirely contained within it. Blocks that belong to multiple clusters are disregarded and the shortest block within each cluster is selected as the representative. (e) The minimum genomic contexts needed to differentiate AMR gene copies are inferred and reads are clustered by the presence of these genes. Full-length paths in node-space are determined by joining the external and internal blocks that share at least one read, then converted to gene-space paths. For each gene-space path, the shortest sub-path that differentiates this path from all others is chosen as the minimum-length path. Reads are clustered based on the presence of the minimum-length paths.

### Estimating the cellular copy number of each AMR gene copy

The cellular copy number of each genomic copy of every previously identified AMR gene is estimated using relative *k*-mer counts, to account for multi-copy plasmids with identical gene content. The frequencies of all *k*-mers (with *k* = 15) in the sample’s FASTQ file are computed using Jellyfish v2.3.0 [39] and a probabilistic mixture model proposed by [40] is used to identify a cut-off separating erroneous from true *k*-mers. Briefly, a log-likelihood function is defined over the *k*-mer frequency spectrum, modelling the probability of each observed count *i* as:

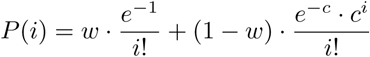

where *w* is the proportion of erroneous *k*-mers, following a Poisson distribution with mean *λ* = 1, and 1 *→ w* is the proportion of correct *k*-mers, assumed to follow a Poisson distribution with mean *λ* = *c*. The parameters *w* and *c* are optimized using the BFGS algorithm implemented in Scipy v1.12.0 [41]. The *k*-mer count cut-off is then selected as the smallest integer *i* where the probability of a *k*-mer originating from the fraction of correct *k*-mers is greater than the error fraction. All *k*-mers with counts below this cut-off are discarded. Next, the filtered *k*-mer count histogram is smoothed using a Savitzky–Golay filter (window length = 30, polynomial order = 3) and the mean read depth is estimated as the count corresponding to the highest peak in the smoothed histogram. Finally, for the minimum-length path of each genomic copy of the AMR genes identified earlier, the filtered histogram is used to compute the median *k*-mer count across all *k*-mers that intersect with the path. This value is divided by the product of the estimated mean read depth and the number of copies of the specific AMR gene within the path to obtain an estimate for the cellular copy number.

### Obtaining AMR gene alleles

The final step is to obtain the allelic sequences of each AMR gene genomic copy. Minimap2 v2.17 [42] is used to map the FASTQ file of the nucleotide sequences of each AMR gene copy to all reference alleles that are contained in the cluster of orthologous genes in the panRG for that AMR gene. The reference allele that most closely matches the reads in the FASTQ with identity *↓*90% and coverage *↓*90% by default is selected, and five iterations of polishing using racon v1.5.0 [17] is conducted to obtain the final sequence of each AMR gene copy.

### Evaluating on simulated data

We developed a snakemake [29] workflow to simulate *E. coli* samples with varying complexities in AMR gene content to assess the accuracy of Amira gene calling and its correlation with read depth and length. A synthetic assembly was generated using the *E. coli K-12* reference sequence (Accession *NC 000913.3*) after removing a single copy of *blaEC* at positions 4377811-4378944, and an *ATCC 11775* reference plasmid (Accession *CP033091.2*) that contains no AMR genes. The workflow inserts complex AMR regions identified from the literature into the synthetic assembly, then simulates sequencing read sets for input into Amira using badread v0.4.1 [20], with parameters `--error model Nanopore2023 and all combinations of: --quantity 10x, 20x, 40x, 80x and --length 5000,6615.5, 10000,10815.5, 20000,19215.5, 40000,36015.5`. Reads were simulated using just the backbone chromosome and plasmid as a negative control, and five additional test cases were simulated using increasingly complex AMR gene containing regions that were identified from public datasets. Genomic-copy-number recall was defined as the mean, across all genes, of the proportion of true genomic copies correctly identified for each method, calculated as the minimum of the true and predicted genomic copy number, divided by the true genomic copy number. We compared the genomic-copy-number recall of Amira v0.9.3 to Flye v2.9.3 with AMRFinderPlus v3.12.8 [7] with database v2024-01-31.1 and options `--plus --organism Escherichia`. Genes with “PARTIALX” or “PARTIALP” in the “Method” column of the AMRFinderPlus output were excluded.

### Evaluating on empirical data

The Nanopore and Illumina reads for 32 *E.coli* isolates in supplementary table S1 were used in the empirical evaluation. Ten of the samples were sequenced using R10.4.1 flow cells (referred to as the R10.4.1 samples) and 22 using R9.4.1 flow cells (referred to as the R9.4.1 samples). The publicly available RefSeq assemblies for the R9.4.1 samples were also retrieved to define the true AMR gene content for these samples [43].

To sequence the ten R10.4.1 samples, isolates were streaked from a single colony onto horse blood agar and cultured overnight at 37°C. DNA was extracted from bacteria using Qiagen DNeasy blood and tissue kit (69504). ONT libraries were prepared using the Rapid barcoding kit (SQK RBK114.96) and sequenced on a PromethION flow cell (FLO-PRO114M) on a PromethION 2 Solo. All data was basecalled using Dorado v0.6.2 [44] and the sup@v4.3.0 model. We retrieved the publicly available Illumina reads for these samples as they had been generated in previous project [45] (Accession PRJNA529744). The long reads were filtered with Filtlong v0.2.1 (--min length 1000) and then assembled using Trycycler v0.5.5 following the ”extra-thorough” assembly instructions in the Trycycler documentation. The resulting consensus assemblies were polished with Illumina data using Polypolish v0.6.0 followed by Pypolca v0.3.1 [46]. A Unicycler v0.5.1 hybrid assembly was also performed to check for small plasmids that may have been missed.

The Illumina and Nanopore reads were mapped to the Trycycler and RefSeq assemblies using minimap2 v2.28 [42] with options `--eqx -a -x sr --secondary=no for the Illumina reads, and --eqx -a -x map-ont --secondary=no` for the Nanopore reads. The Nanopore read depths across each assembly were visualized, normalized by the mean read depth for each contig containing an AMR gene, to check for regions of elevated coverage that may indicate collapsed duplications, that were present in two samples. The reads mapping to the problematic contigs were subsetted with samtools v1.17 [47] and pyfastaq v3.17.0 [48] and reassembled using Canu v2.3 [49] with genomeSize=20k. One round of short-read polishing was then conduced for each with Polypolish v0.6.0 [46]. TNA v0.3.0 [50] was used to visualize the alignments of the reassembled contigs to the reference, and the new assembly was only accepted if the alignment had no further signal of duplication or any other problem. Further information detailing the assembly validation be found in figures S2-S7.

A true genomic copy of an AMR gene in the truth assemblies is defined as any sequence present in the assembly that is at least 90% identical to and 90% covered by a reference allele of an AMR gene that is included in the panRG. This was done by mapping all the AMR gene reference alleles included in the reference panRG to each assembly using minimap2. All hits with alignment coverage *<*90% and identity *<*90% were filtered and the remaining hits were sorted first by identity, then by length. The best match was chosen in cases where multiple alleles overlapped with each other. False pseudogenes are frequent in bacterial assemblies [51] and this relaxed coverage threshold avoided penalization of AMR genes that were present in the truth assemblies but contained base-level errors leading to frameshifts or premature stop codons within open reading frames. All of the tools tested call AMR genes that are less than 100% covered by a reference.

The Nanopore read set for each sample was subsampled to 200x using rasusa v2.1.0 [52] and the accuracy of Amira v0.9.3 was compared to that of AMRFinderPlus v3.12.8 [7], with database v2024-01-31.1 and options `--plus --organism` Escherichia, run on Nanopore-only assemblies generated with Flye v2.9.3 [21] and Raven v1.8.3 [22], and hybrid assemblies generated with Unicycler v0.5.0 [23]. All assemblers were run using their default parameters and genes with “PARTIALX” or “PARTIALP” in the “Method” column of the AMRFinderPlus output were excluded. Resfinder [8] v4.6.0 was also ran on the Nanopore reads for each sample with options `-s e.coli -acq --Nanopore` by generating an index of all AMR alleles in our reference panRG using kma index v1.4.15 [53]. All ResFinder hits with *<*90% identity or *<*90% coverage were excluded. For each flow cell type, genomic-copy-number recall was defined as the average, across all samples, of the proportion of true gene copies correctly identified by each method (calculated as the minimum of the true and predicted copy numbers divided by the true copy number). Precision was defined as the average proportion of predicted genomic copies that were true genomic copies (calculated as the maximum of zero and the difference between the predicted and true genomic copy numbers, divided by the true genomic copy number). The true cellular copy numbers of each genomic copy of an AMR gene were estimated by obtaining the mean Nanopore read depth across the contig for the gene, normalized by the mean read depth across the longest contig in each sample. All alleles for each unique AMR gene present in both the reference assembly and the output of the evaluated tool were aligned using MAFFT v7.526 --auto [54]. Nucleotide-level accuracies were calculated as the proportion of matching columns in the alignment between each reference allele and the output allele. Each output allele was matched to the closest reference allele without replacement, prioritizing nucleotide accuracy followed by the absolute difference in cellular copy number.

Divergent paths in a single connected component of the Amira graph were resulting in false positive gene calls in one sample (identifier AUSMDU00021208). By subsetting the reads unique to the differing paths and visualizing the pileups with IGV v2.19.1 [55], this was found to be the result of the presence of a minor frequency structural variants of a plasmid in this sample (Fig.S9). We chose not to penalize the assemblers for missing the low-frequency variant, nor penalize Amira for identifying it as it resulted from genuine heterogeneity present in the read sets.

### Estimating AMR gene frequencies across large datasets of long read sequences

We used Amira to analyze 8580 *E. coli*, 2448 *K. pneumoniae* and 415 *E. faecium* samples in the ENA with long reads available. A TSV of all run accessions for Tax Id 562 for *E. coli*, Tax Id 563 for *K. pneumoniae* and Tax Id 1352 for *E. faecium* (as of 11 February 2025) was obtained from the ENA, and all read sets downloaded if they contained the terms “Nanopore”, “MinION”, “PromethION” or “PacBio” in the “description column of the TSVs. Amira *v0.9.3* was then run with option `-species Escherichia coli, Klebsiella pneumoniae or Enterococcus faecium, as was AMRFinderPlus v3.12.8 with options -plus -organism Escherichia, Klebsiella pneumoniae or Enterococcus faecium` on long read-only assemblies generated with Flye v2.9.3 with default parameters. For the union of the AMR genes identified by either Amira or Flye AMRFP, we calculated the relative frequency of the presence of the gene for each species. We verified that each gene called by Amira was supported by the reads (and not the result of overcalling) by mapping the long read FASTQ for each sample to the nucleotide sequence for the gene that was assembled by Amira, ensuring *↓*85% alignment coverage across the sequence using samtools v1.17 coverage. Samples where Amira and Flye did not complete within 240 minutes, samples that failed to run for either tool and genes with a value of *<*85% in the “% Coverage of reference sequence” column of the AMRFinderPlus output were excluded.

## Declarations

### Ethics approval and consent to participate

Not applicable.

### Consent for publication

Not applicable.

### Availability of data and materials

Amira is freely available from https://github.com/Danderson123/Amira under an Apache-2.0 license and can be installed via PyPI, Conda [56] or Singularity [57]. The panRG construction pipeline is available from https://github.com/Danderson123/Amira_panRG_pipeline. Snakemake workflows to rerun the analyses in this paper and the long read accessions for the samples used in the ENA evaluation are available at https://github.com/Danderson123/amira_paper. The *E. coli* panRG is available from [58], the *K. pneumoniae* panRG is available from [59], the *E. faecium* panRG from [60] the *S. pneumoniae* panRG from [61] and the *S. aureus* panRG from [62]. The reference assemblies for all of the empirical evaluation samples are available from [63] and the long and short read accessions can be found in table S1.

### Authors’ contributions

Conceptualization: D.A., Z.I. Data Curation: D.A, R.W., T.L., L.J. Formal analysis: D.A. Funding acquisition: Z.I. Investigation: D.A. Methodology: D.A., L.L, Z.I. Project administration: Z.I. Resources: L.J., R.W., Z.I. Software: D.A. Supervision: Z.I. Validation: D.A. Visualisation: D.A. Writing – original draft: D.A., R.W., T.L., Z.I. Writing – review & editing: D.A., R.W., Z.I.

## Acknowledgements

The authors are very grateful to Martin Hunt, Leah Roberts and Wei Shen for critical review of the manuscript, William Matlock for discussions about MIC prediction, and John Lees, Julian Parkhill and Nassos Typas for their valuable advice, guidance and support throughout the project as part of the thesis advisory committee for DA. We also thank Daria Frolova and Samuel Horsfield for their general interest and providing fruitful discussion during the undertaking of this work. This paper also acknowledges the PulseNet Asia-Pacific team at the Centre for Pathogen Genomics for contributing data to the study.

## References

[1] Naghavi M, Vollset SE, Ikuta KS, Swetschinski LR, Gray AP, Wool EE, et al. Global burden of bacterial antimicrobial resistance 1990–2021: a systematic analysis with forecasts to 2050. The Lancet. 2024;404(10459):1199–1226. 10.1016/S0140-6736(24)01867-1.

[2] Zhao S, Tyson GH, Chen Y, Li C, Mukherjee S, Young S, et al. Whole-Genome Sequencing Analysis Accurately Predicts Antimicrobial Resistance Phenotypes in Campylobacter spp. Applied and Environmental Microbiology. 2016;82(2):459–466. 10.1128/AEM.02873-15. https://journals.asm.org/doi/pdf/10.1128/aem.02873-15.

[3] Davies TJ, Stoesser N, Sheppard AE, Abuoun M, Fowler P, Swann J, et al. Reconciling the Potentially Irreconcilable? Genotypic and Phenotypic Amoxicillin-Clavulanate Resistance in Escherichia coli. Antimicrobial Agents and Chemotherapy. 2020;64(6):e02026–19. 10.1128/aac.02026-19.

[4] Marshall SH, Donskey CJ, Hutton-Thomas R, Salata RA, Rice LB. Gene dosage and linezolid resistance in Enterococcus faecium and Enterococcus faecalis [Journal Article]. Antimicrobial Agents and Chemotherapy. 2002 Oct;46(10):3334– 3336. 10.1128/AAC.46.10.3334-3336.2002.

[5] Partridge SR, Kwong SM, Firth N, Jensen SO. Mobile Genetic Elements Associated with Antimicrobial Resistance. Clinical Microbiology Reviews. 2018 October;31(4):e00088–17. 10.1128/CMR.00088-17.

[6] Koren S, Phillippy AM. One chromosome, one contig: complete microbial genomes from long-read sequencing and assembly. Current Opinion in Microbiology. 2015;23:110–120. Host–microbe interactions: bacteria • Genomics. 10.1016/j.mib.2014.11.014.

[7] Feldgarden M, Brover V, Gonzalez-Escalona N, Frye JG, Haendiges J, Haft DH, et al. AMRFinderPlus and the Reference Gene Catalog facilitate examination of the genomic links among antimicrobial resistance, stress response, and virulence. Scientific Reports. 2021 Jun;11(1):12728. 10.1038/s41598-021-91456-0.

[8] Bortolaia V, Kaas RS, Ruppe E, Roberts MC, Schwarz S, Cattoir V, et al. ResFinder 4.0 for predictions of phenotypes from genotypes. Journal of Antimicrobial Chemotherapy. 2020 December;75(12):3491–3500. 10.1093/jac/dkaa345.

[9] Alcock BP, Raphenya AR, Lau TTY, Tsang KK, Bouchard M, Edalatmand A, et al. CARD 2020: antibiotic resistome surveillance with the comprehensive antibiotic resistance database. Nucleic Acids Research. 2019;48(D1):D517–D525. 10.1093/nar/gkz935.

[10] Seemann T.: Abricate. GitHub repository. https://github.com/tseemann/abricate.

[11] Hunt M, Mather AE, Sánchez-Busó L, Page AJ, Parkhill J, Keane JA, et al. ARIBA: rapid antimicrobial resistance genotyping directly from sequencing reads. Microbial Genomics. 2017 Sep;3(10):e000131. 10.1099/mgen.0.000131.

[12] Bradley P, Gordon NC, Walker TM, Dunn L, Heys S, Huang B, et al. Rapid antibiotic-resistance predictions from genome sequence data for *Staphylococcus aureus* and *Mycobacterium tuberculosis*. Nature Communications. 2015 Dec;6:10063. 10.1038/ncomms10063.

[13] Wick RR, Holt KE. Benchmarking of long-read assemblers for prokaryote whole genome sequencing [version 1; peer review: 4 approved]. F1000Research. 2019;8:2138. 10.12688/f1000research.21782.1.

[14] Wick RR, Judd LM, Cerdeira LT, Hawkey J, Méric G, Vezina B, et al. Trycycler: consensus long-read assemblies for bacterial genomes. Genome Biology. 2021;22(1):266. 10.1186/s13059-021-02483-z.

[15] Foster-Nyarko E, Cottingham H, Wick RR, Judd LM, Lam MMC, Wyres KL, et al. Nanopore-only assemblies for genomic surveillance of the global priority drug-resistant pathogen, Klebsiella pneumoniae. Microbial Genomics. 2023;9(2):mgen000936. 10.1099/mgen.0.000936.

[16] Colquhoun RM, Hall MB, Lima L, Roberts LW, Malone KM, Hunt M, et al. Pandora: nucleotide-resolution bacterial pan-genomics with reference graphs. Genome Biology. 2021;22(1):267. 10.1186/s13059-021-02473-1.

[17] Vaser R, Sović I, Nagarajan N, Šikić M. Fast and accurate de novo genome assembly from long uncorrected reads. Genome Research. 2017 May;27(5):737– 746. 10.1101/gr.214270.116.

[18] Robertson J, Nash JHE. MOB-suite: software tools for clustering, reconstruction and typing of plasmids from draft assemblies [Journal Article]. Microbial Genomics. 2018;4(8). 10.1099/mgen.0.000206.

[19] Tamamura-Andoh Y, Niwa H, Kinoshita Y, Uchida-Fujii E, Arai N, Watanabe-Yanai A, et al. Duplication of blaCTX-M-1 and a class 1 integron on the chromosome enhances antimicrobial resistance in Escherichia coli isolated from racehorses in Japan. Journal of Global Antimicrobial Resistance. 2021 Dec;27:225–227. 10.1016/j.jgar.2021.10.004.

[20] Wick RR. Badread: simulation of error-prone long reads. Journal of Open Source Software. 2019;4(36):1316. 10.21105/joss.01316.

[21] Kolmogorov M, Yuan J, Lin Y, Pevzner PA. Assembly of long, error-prone reads using repeat graphs. Nature Biotechnology. 2019 May;37(5):540–546. 10.1038/s41587-019-0072-8.

[22] Vaser R, Šikić M. Time- and memory-efficient genome assembly with Raven. Nature Computational Science. 2021;1(5):332–336. 10.1038/s43588-021-00073-4.

[23] Wick RR, Judd LM, Gorrie CL, Holt KE. Unicycler: Resolving bacterial genome assemblies from short and long sequencing reads. PLoS Computational Biology. 2017 Jun;13(6):e1005595. 10.1371/journal.pcbi.1005595.

[24] Gorrie CL, Silva AGD, Ingle DJ, Higgs C, Seemann T, Stinear TP, et al. Key parameters for genomics-based real-time detection and tracking of multidrug-resistant bacteria: a systematic analysis. The Lancet Microbe. 2021;2(11):e575– e583. 10.1016/s2666-5247(21)00149-x.

[25] Tonkin-Hill G, MacAlasdair N, Ruis C, Weimann A, Horesh G, Lees JA, et al. Producing polished prokaryotic pangenomes with the Panaroo pipeline. Genome Biology. 2020 jul;21(1):180. 10.1186/s13059-020-02090-4.

[26] Schmartz GP, Hartung A, Hirsch P, Kern F, Fehlmann T, Müller R, et al. PLSDB: advancing a comprehensive database of bacterial plasmids. Nucleic Acids Research. 2021 11;50(D1):D273–D278. 10.1093/nar/gkab1111. https://academic.oup.com/nar/article-pdf/50/D1/D273/42058178/gkab1111.pdf.

[27] Ruan J, Li H. Fast and accurate long-read assembly with wtdbg2. Nature Methods. 2020;17(2):155–158. 10.1038/s41592-019-0669-3.

[28] Matlock W, Rodger G, Pritchard E, Colpus M, Kapel N, Barrett L, et al. E. coli phylogeny drives co-amoxiclav resistance through variable expression of blaTEM-1. bioRxiv. 2024;10.1101/2024.08.12.607562. https://www.biorxiv.org/content/early/2024/08/12/2024.08.12.607562.full.pdf.

[29] Köster J, Rahmann S. Snakemake—a scalable bioinformatics workflow engine. Bioinformatics. 2012 08;28(19):2520–2522. 10.1093/bioinformatics/bts480. https://academic.oup.com/bioinformatics/article-pdf/28/19/2520/48879301/bioinformatics_28_19_2520.pdf.

[30] Schwengers O, Jelonek L, Dieckmann MA, Beyvers S, Blom J, Goesmann A. Bakta: rapid and standardized annotation of bacterial genomes via alignment-free sequence identification. Microb Genom. 2021 nov;7(11):000685. 10.1099/mgen.0.000685.

[31] Bouras G, Houtak G, Wick RR, Mallawaarachchi V, Roach MJ, Papudeshi B, et al. Hybracter: enabling scalable, automated, complete and accurate bacterial genome assemblies. Microbial Genomics. 2024 May;10(5):001244. 10.1099/mgen.0.001244.

[32] Hetland MAK, Winkler MA, Kaspersen H, Håkonsholm F, Bakksjø RJ, Bernhoff E, et al. A genome-wide One Health study of Klebsiella pneumoniae in Norway reveals overlapping populations but few recent transmission events across reservoirs. bioRxiv. 2024;10.1101/2024.09.11.612360. https://www.biorxiv.org/content/early/2024/09/11/2024.09.11.612360.full.pdf.

[33] Li H, Durbin R. Fast and accurate short read alignment with Burrows–Wheeler transform. Bioinformatics. 2009 05;25(14):1754–1760. 10.1093/bioinformatics/btp324. https://academic.oup.com/bioinformatics/article-pdf/25/14/1754/48994219/bioinformatics_25_14_1754.pdf.

[34] Danecek P, Bonfield JK, Liddle J, Marshall J, Ohan V, Pollard MO, et al. Twelve years of SAMtools and BCFtools. GigaScience. 2021 02;10(2):giab008. 10.1093/gigascience/giab008. https://academic.oup.com/gigascience/article-pdf/10/2/giab008/60687743/giab008.pdf.

[35] Price MN, Dehal PS, Arkin AP. FastTree 2 – Approximately Maximum-Likelihood Trees for Large Alignments. PLOS ONE. 2010 03;5(3):1–10. 10.1371/journal.pone.0009490.

[36] Tonkin-Hill G, Lees JA, Bentley SD, Frost SDW, Corander J. Fast hierarchical Bayesian analysis of population structure. Nucleic Acids Research. 2019 05;47(11):5539–5549. 10.1093/nar/gkz361. https://academic.oup.com/nar/article-pdf/47/11/5539/28839395/gkz361.pdf.

[37] Arredondo-Alonso S, Top J, Schürch A, McNally A, Puranen S, Pesonen M, et al. Genomes of a major nosocomial pathogen Enterococcus faecium are shaped by adaptive evolution of the chromosome and plasmidome. bioRxiv. 2019;10.1101/530725. https://www.biorxiv.org/content/early/2019/01/28/530725.full.pdf.

[38] Pierce NT, Irber L, Reiter T, Brooks P, Brown CT. Large-scale sequence comparisons with sourmash. F1000Research. 2019 Jul;8:1006. 10.12688/f1000research.19675.1.

[39] Marçais G, Kingsford C. A fast, lock-free approach for efficient parallel counting of occurrences of k-mers. Bioinformatics. 2011;27(6):764–770. 10.1093/bioinformatics/btr011.

[40] Derelle R, Wachsmann Jv, Máklin T, Hellewell J, Russell T, Lalvani A, et al. Seamless, rapid, and accurate analyses of outbreak genomic data using split k-mer analysis. Genome Research. 2024;34(10):1661–1673. 10.1101/gr.279449.124.

[41] Virtanen P, Gommers R, Oliphant TE, Haberland M, Reddy T, Cournapeau D, et al. SciPy 1.0: Fundamental Algorithms for Scientific Computing in Python. Nature Methods. 2020;17:261–272. 10.1038/s41592-019-0686-2.

[42] Li H. Minimap2: pairwise alignment for nucleotide sequences. Bioinformatics. 2018 05;34(18):3094–3100. 10.1093/bioinformatics/bty191. https://academic.oup.com/bioinformatics/article-pdf/34/18/3094/48919122/bioinformatics_34_18_3094.pdf.

[43] Moran RA, Baomo L, Doughty EL, Guo Y, Ba X, van Schaik W, et al. Extended-Spectrum-Lactamase Genes Traverse the Escherichia coli Populations of Intensive Care Unit Patients, Staff, and Environment. Microbiology Spectrum. 2023;11(2):e05074–22. 10.1128/spectrum.05074-22. https://journals.asm.org/doi/pdf/10.1128/spectrum.05074-22.

[44] PLC ON.: Dorado. GitHub. https://github.com/nanoporetech/dorado.

[45] Sherry NL, Lane CR, Kwong JC, Schultz M, Sait M, Stevens K, et al. Genomics for Molecular Epidemiology and Detecting Transmission of Carbapenemase-Producing ¡i¿Enterobacterales¡/i¿ in Victoria, Australia, 2012 to 2016. Journal of Clinical Microbiology. 2019;57(9):10.1128/jcm.00573–19. 10. 1128/jcm.00573-19. https://journals.asm.org/doi/pdf/10.1128/jcm.00573-19.

[46] Bouras G, Judd LM, Edwards RA, Vreugde S, Stinear TP, Wick RR. How low can you go? Short-read polishing of Oxford Nanopore bacterial genome assemblies [Journal Article]. Microbial Genomics. 2024;10(6). 10.1099/mgen.0.001254.

[47] Li H, Handsaker B, Wysoker A, Fennell T, Ruan J, Homer N, et al. The Sequence Alignment/Map format and SAMtools. Bioinformatics. 2009 aug;25(16):2078– 2079. 10.1093/bioinformatics/btp352.

[48] Sanger-Pathogens.: Fastaq. GitHub repository. https://github.com/sanger-pathogens/Fastaq.

[49] Koren S, Walenz BP, Berlin K, Miller JR, Bergman NH, Phillippy AM. Canu: scalable and accurate long-read assembly via adaptive k-mer weighting and repeat separation. Genome Research. 2017 may;27(5):722–736. 10.1101/gr.215087.116.

[50] Hunt M.: TNA. GitHub repository. https://github.com/martinghunt/tna.

[51] Cooley NP, Wright ES. Many purported pseudogenes in bacterial genomes are bona fide genes. BMC Genomics. 2024 apr;25(1):365. 10.1186/s12864-024-10137-0.

[52] Hall MB. Rasusa: Randomly subsample sequencing reads to a specified coverage. Journal of Open Source Software. 2022;7(69):3941. 10.21105/joss. 03941.

[53] Clausen PTLC, Aarestrup FM, Lund O. Rapid and precise alignment of raw reads against redundant databases with KMA. BMC Bioinformatics. 2018 Aug;19(1):307. 10.1186/s12859-018-2336-6.

[54] Katoh K, Misawa K, Kuma Ki, Miyata T. MAFFT: a novel method for rapid multiple sequence alignment based on fast Fourier transform. Nucleic Acids Research. 2002 Jul;30(14):3059–3066. 10.1093/nar/gkf436.

[55] Thorvaldsdóttir H, Robinson JT, Mesirov JP. Integrative Genomics Viewer (IGV): high-performance genomics data visualization and exploration. Briefings in Bioinformatics. 2012 04;14(2):178–192. 10.1093/bib/bbs017. https://academic.oup.com/bib/article-pdf/14/2/178/546734/bbs017.pdf.

[56] Grüning B, Dale R, Sjödin A, Chapman BA, Rowe J, Tomkins-Tinch CH, et al. Bioconda: sustainable and comprehensive software distribution for the life sciences. Nature Methods. 2018 Jul;15(7):475–476. 10.1038/s41592-018-0046-7.

[57] Kurtzer GM, Sochat V, Bauer MW. Singularity: Scientific containers for mobility of compute. PLOS ONE. 2017 05;12(5):1–20. 10.1371/journal.pone.0177459.

[58] Anderson D. Escherichia coli panRG. 2025 4;10.6084/m9.figshare.28723748.v2.

[59] Anderson D. Klebsiella pneumoniae panRG. 2025 4;10.6084/m9.figshare.28723775.v1.

[60] Anderson D. Enterococcus faecium panRG. 2025 4;10.6084/m9.figshare.28723262.v1.

[61] Anderson D. Streptococcus pneumoniae panRG. 2025 4;10.6084/m9.figshare.28768637.v1.

[62] Anderson D. Staphylococcus aureus panRG. 2025 4;10.6084/m9.figshare.28768619.v1.

[63] Anderson D. Reference assemblies used in the empirical evaluation in the Amira paper. 2025 5;10.6084/m9.figshare.28958849.v1.

[64] Katz LS, Griswold T, Morrison SS, Caravas JA, Zhang S, den Bakker HC, et al. Mashtree: a rapid comparison of whole genome sequence files. Journal of Open Source Software. 2019;4(44):1762. 10.21105/joss.01762.

