## Supplementary Figures and Tables for "Amira: detection of AMR genes directly from long reads using gene-space *de Bruijn* graphs"

### Supplemental information

**Table S1** Nanopore and Illumina accession, Nanopore flow cell technology and identifier for the 32 *E. coli* used in the truth evaluation.

| Sample name | Nanopore accession | Nanopore flow cell | Illumina accession |
|---|---|---|---|
| GCA_027944575.1_ASM2794457v1_genomic | SRR23044210 | R9.4.1 | SRR22543847 |
| GCA_027944595.1_ASM2794459v1_genomic | SRR23044211 | R9.4.1 | SRR22543854 |
| GCA_027944615.1_ASM2794461v1_genomic | SRR23044212 | R9.4.1 | SRR22543861 |
| GCA_027944635.1_ASM2794463v1_genomic | SRR23044213 | R9.4.1 | SRR22543882 |
| GCA_027944655.1_ASM2794465v1_genomic | SRR23044215 | R9.4.1 | SRR22543883 |
| GCA_027944675.1_ASM2794467v1_genomic | SRR23044216 | R9.4.1 | SRR22543887 |
| GCA_027944695.1_ASM2794469v1_genomic | SRR23044217 | R9.4.1 | SRR22543892 |
| GCA_027944715.1_ASM2794471v1_genomic | SRR23044218 | R9.4.1 | SRR22543930 |
| GCA_027944735.1_ASM2794473v1_genomic | SRR23044219 | R9.4.1 | SRR22543931 |
| GCA_027944775.1_ASM2794477v1_genomic | SRR23044221 | R9.4.1 | SRR22543936 |
| GCA_027944795.1_ASM2794479v1_genomic | SRR23044222 | R9.4.1 | SRR22543941 |
| GCA_027944815.1_ASM2794481v1_genomic | SRR23044223 | R9.4.1 | SRR22543945 |
| GCA_027944835.1_ASM2794483v1_genomic | SRR23044224 | R9.4.1 | SRR22543947 |
| GCA_027944875.1_ASM2794487v1_genomic | SRR23044204 | R9.4.1 | SRR22543793 |
| GCA_027944895.1_ASM2794489v1_genomic | SRR23044205 | R9.4.1 | SRR22543798 |
| GCA_027944915.1_ASM2794491v1_genomic | SRR23044206 | R9.4.1 | SRR22543803 |
| GCA_027944935.1_ASM2794493v1_genomic | SRR23044207 | R9.4.1 | SRR22543900 |
| GCA_027944955.1_ASM2794495v1_genomic | SRR23044208 | R9.4.1 | SRR22543904 |
| GCA_027945015.1_ASM2794501v1_genomic | SRR23044214 | R9.4.1 | SRR22543922 |
| GCA_027945035.1_ASM2794503v1_genomic | SRR23044225 | R9.4.1 | SRR22543827 |
| GCA_027945055.1_ASM2794505v1_genomic | SRR23044226 | R9.4.1 | SRR22543837 |
| GCA_028551585.1_ASM2855158v1_genomic | SRR23044220 | R9.4.1 | SRR22543934 |
| AUSMDU00010405 | SRR32405434 | R10.4.1 | SRR15116130 |
| AUSMDU00015264 | SRR32405436 | R10.4.1 | SRR14673594 |
| AUSMDU00021208 | SRR32405439 | R10.4.1 | SRR15116365 |
| AUSMDU00031899 | SRR32405433 | R10.4.1 | SRR15116275 |
| AUSMDU00031978 | SRR32405440 | R10.4.1 | SRR15116274 |
| AUSMDU00032793 | SRR32405441 | R10.4.1 | SRR15116259 |
| AUSMDU00036400 | SRR32405435 | R10.4.1 | SRR15116221 |
| AUSMDU00040126 | SRR32405437 | R10.4.1 | SRR1511617 |
| AUSMDU00055259 | SRR32405442 | R10.4.1 | SRR28786155 |
| AUSMDU00062512 | SRR32405438 | R10.4.1 | SRR32390598 |

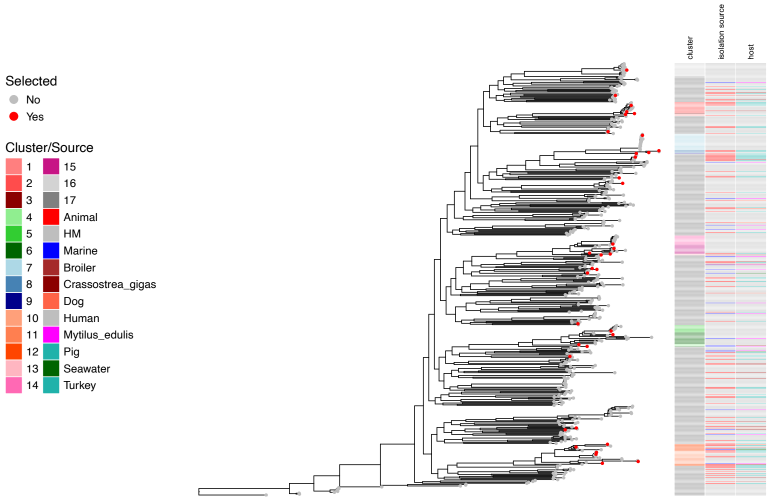


**Fig. S1** Heatmap displaying the *Klebsiella pneumoniae* phylogenetic tree annotated with clusters identified by FastBAPS and the isolation source. Red tips indicate the isolates that were selected for panRG construction.

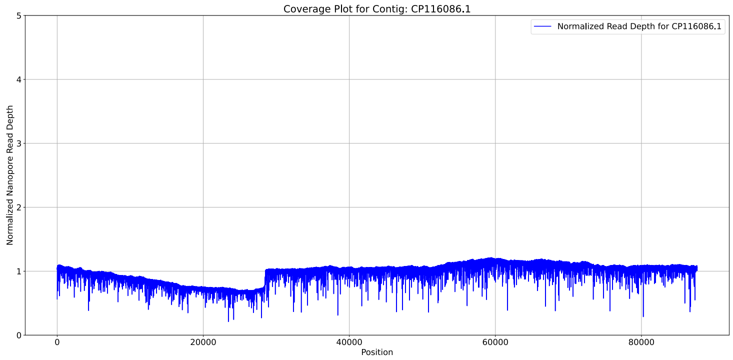


**Fig. S2** We identified a region of elevated coverage in contig CP116086.1 in sample GCA_027944635.1_ASM2794463v1_genomic. We extracted the reads mapping to this region using `samtools v1.17 view GCA_027944635.1_ASM2794463v1_genomic.bam CP116086.1 | cut -f1 | sort | uniq` and subsetted the nanopore FASTQ to just these reads using pyfastaq `v3.17.0 filter`. We reassembled the reads using Canu `v2.3` with `genomeSize=100k` and short-read polished the resulting contig with polypolish `v0.6.0`. We then directly replaced CP116086.1 with this contig.

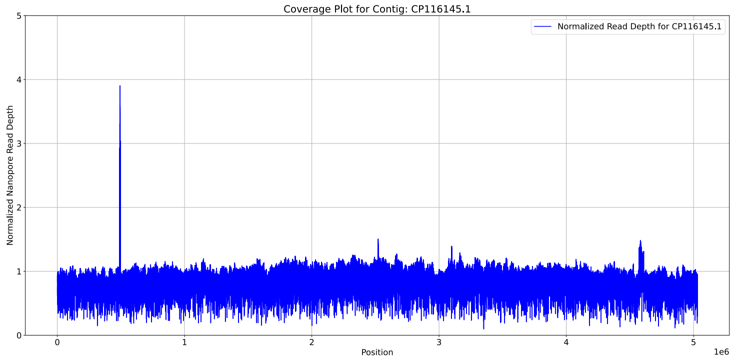


**Fig. S3** We identified a region of elevated coverage in contig CP116145.1 in sample GCA_027944835.1_ASM2794483v1_genomic. We extracted the soft-clipped reads mapping to this region using `samtools v1.17 view -f 2048 -f 2064 GCA_027944835.1_ASM2794483v1_genomic.bam CP116145.1:300000-700000 | cut -f1 | sort | uniq` and subsetted the nanopore FASTQ to just these reads using pyfastaq `v3.17.0 filter`. We reassembled the reads using flye `v2.9.3 --iterations 3`. We visualised the alignment between the assembly and the reference using TNA `v0.3.0`, and found it fully covered plasmid contig CP116148.1 with 99.9% identity so was not evident of a collapsed duplication.

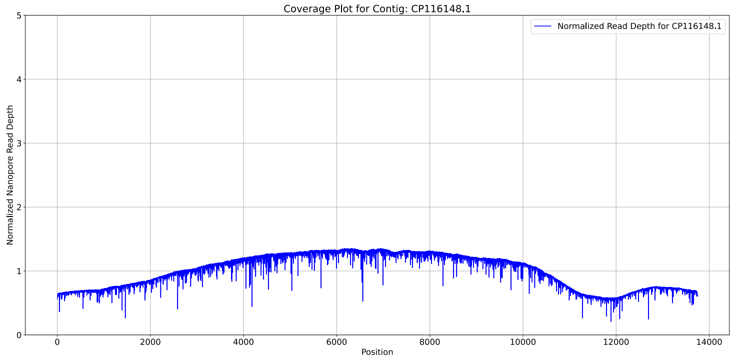


**Fig. S4** We identified a region of elevated coverage in contig CP116148.1 in sample GCA_027944835.1_ASM2794483v1_genomic. We subsetted the reads mapping to the contig using `samtools v1.17 view GCA_027944835.1_ASM2794483v1_genomic.bam CP116148.1 | cut -f1 | sort | uniq` and pyfastaq `v3.17.0 filter`. We reassembled the reads using Canu `v2.3` with `genomeSize=20k` and short-read polished the resulting contig with polypolish `v0.6.0`. We then directly replaced CP116148.1 with this contig.

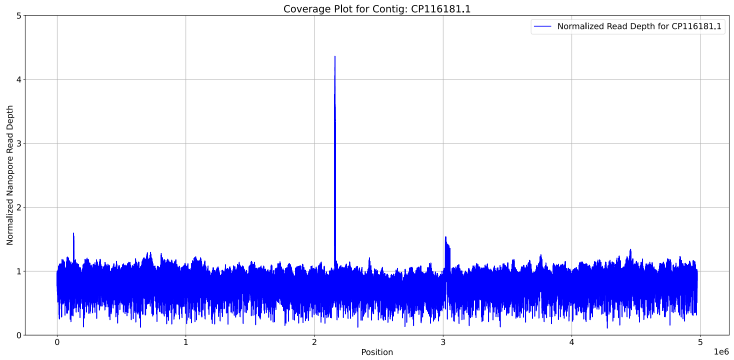


**Fig. S5** We identified a region of elevated coverage in contig CP116181.1 in sample GCA_027944955.1_ASM2794495v1_genomic. We extracted the soft-clipped reads mapping to this region using `samtools v1.17 view -f 2048 -f 2064 GCA_027944955.1_ASM2794495v1_genomic.bam CP116181.1:2000000-2500000 | cut -f1 | sort | uniq` and subsetted the nanopore FASTQ to just these reads using pyfastaq `v3.17.0 filter`. We reassembled the reads using flye `v2.9.3 --iterations 3`. We visualised the alignment between the assembly and the reference using TNA `v0.3.0`, and found it fully covered plasmid contig CP116184.1 with identity 99.8%, so was not evident of a collapsed duplication.

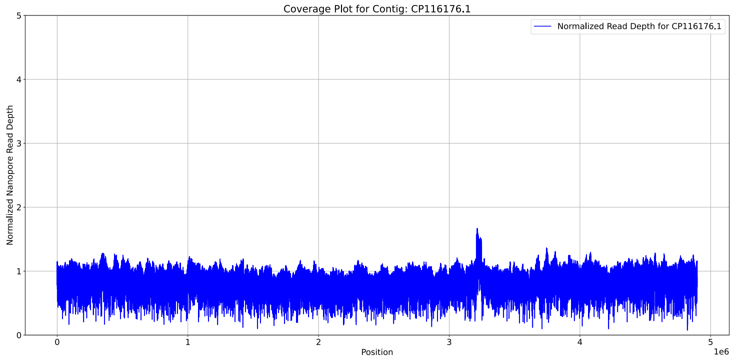


**Fig. S6** We identified a region of elevated coverage in contig CP116176.1 in sample GCA_027944935.1_ASM2794493v1_genomic. We extracted the soft-clipped reads mapping to this region using `samtools v1.17 view -f 2048 -f 2064 sample GCA_027944935.1_ASM2794493v1_genomic.bam CP116176.1:3100000-35000000 | cut -f1 | sort | uniq` and subsetted the nanopore FASTQ to just these reads using pyfastaq `v3.17.0 filter`. We reassembled the reads using flye `v2.9.3 --iterations 3`. We visualised the alignment between the assembly and the reference using TNA `v0.3.0`, and found all generated contigs full aligned to the reference with >95% identity.

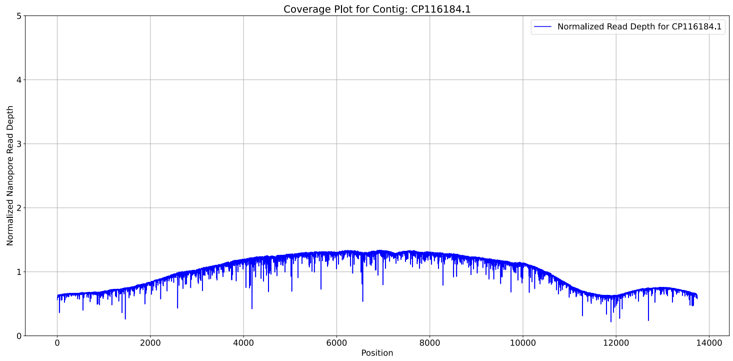


**Fig. S7** We identified a region of elevated coverage in contig CP116184.1 in sample GCA_027944955.1_ASM2794495v1_genomic. We subsetted the reads mapping to the contig using `samtools v1.17 view GCA_027944955.1_ASM2794495v1_genomic.bam CP116184.1 | cut -f1 | sort | uniq` and pyfastaq `v3.17.0 filter`. We reassembled the reads using Canu `v2.3` with `genomeSize=20k` and short-read polished the resulting contigs with polypolish `v0.6.0`. We then directly replaced CP116184.1 with the longest polished contig.

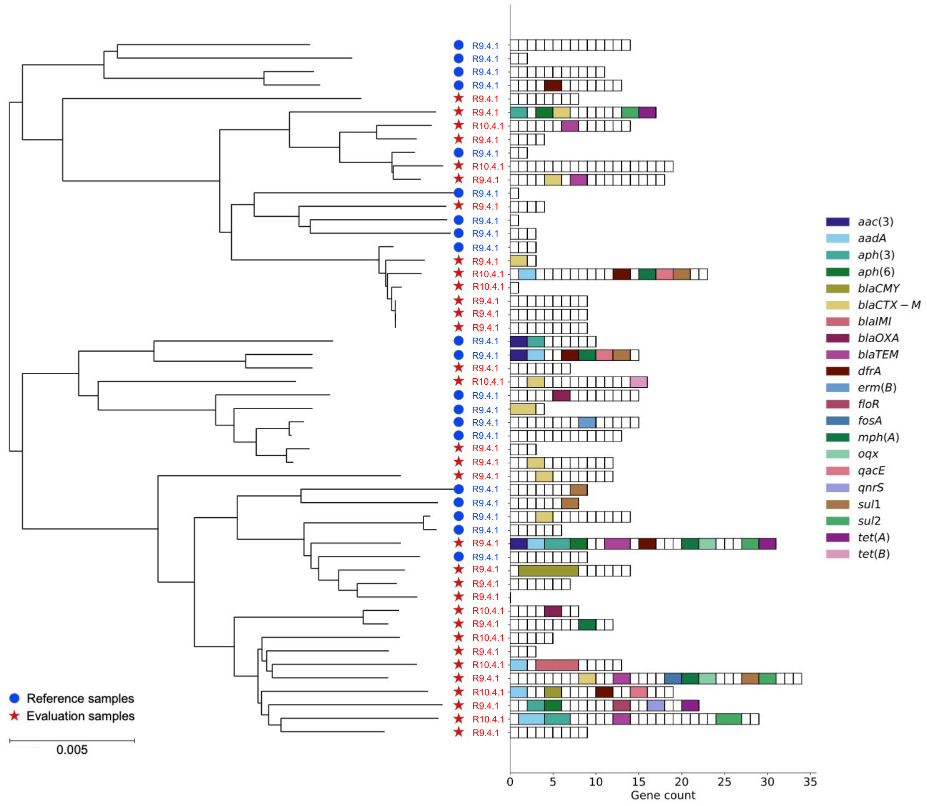

**Fig. S8** A phylogeny showing the relationship between the samples used to construct the reference panRG and the samples used to evaluate the accuracy of Amira, alongside the Nanopore flow cell used for sequencing and the AMR gene content of samples used in the truth evaluation. Right: The x-axis is the number of AMR gene copies, the y-axis is the sample identifier and each block in each bar represents a unique AMR gene. AMR genes with more than one copy in a given sample have been colored. Left: The phylogeny was constructed using mashtree v1.4.6 [64] and the order of samples matches the samples at the tips in the phylogeny.

**Table S2** Benchmarking of the computational resources used by Amira and competing tools in the empirical evaluation of 32 *E. coli* samples. All methods were run using 4 CPUs.

| Method | Range wall-clock time | Range peak RAM |
|---|---|---|
| Amira | 605 - 1692 s | 4.4 - 12.5 GB |
| Flye AMRFP (Flye, AMRFP) | 1071 - 4087 (957 - 4032, 24 - 178) s | 9.8 - 11.4 (9.8 - 11.4, 0.4 - 0.4) GB |
| Raven AMRFP (Raven, AMRFP) | 377 - 1189 (291 - 1107, 48 - 153) s | 3.7 - 12.1 (3.7 - 12.1, 0.4 - 0.4) GB |
| ResFinder | 170 - 692 s | 0.3 - 0.3 GB |
| Unicycler AMRFP (Unicycler, AMRFP) | 6579 - 31983 (6502 - 31882, 65 - 152) s | 3.5 - 9.1 (3.5 - 9.1, 0.4 - 0.4) GB |

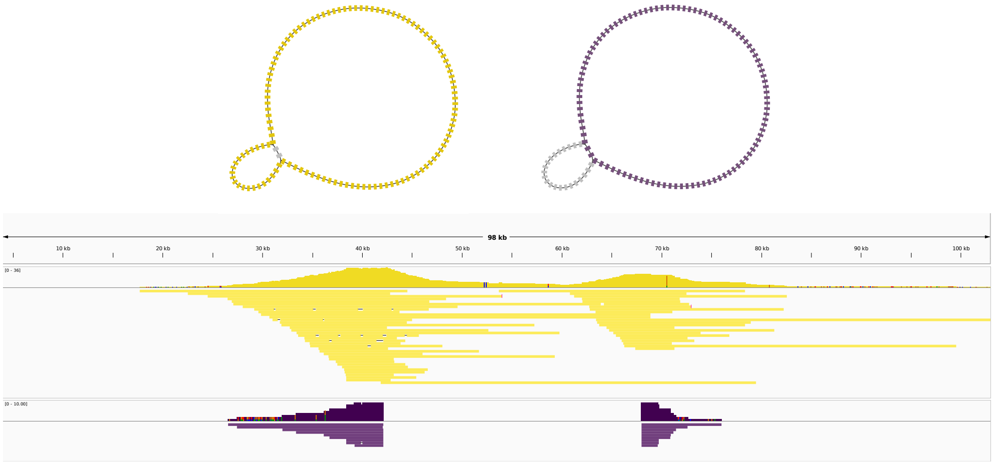


**Fig. S9** A comparison of the reads corresponding to two structural variants of the same plasmid in sample AUSMDU00021208, one version containing a 26.7$kbp$ mobile genetic element (yellow) and one version without it (purple). The variant without the element is missing from the reference assembly for this sample, resulting in Amira calling additional AMR gene copies due to an additional path through the connected component (top). We separated the reads unique to each path, mapped them to the reference and visualised the pileups with IGV to find two variants of the plasmid exist in the read set (bottom)

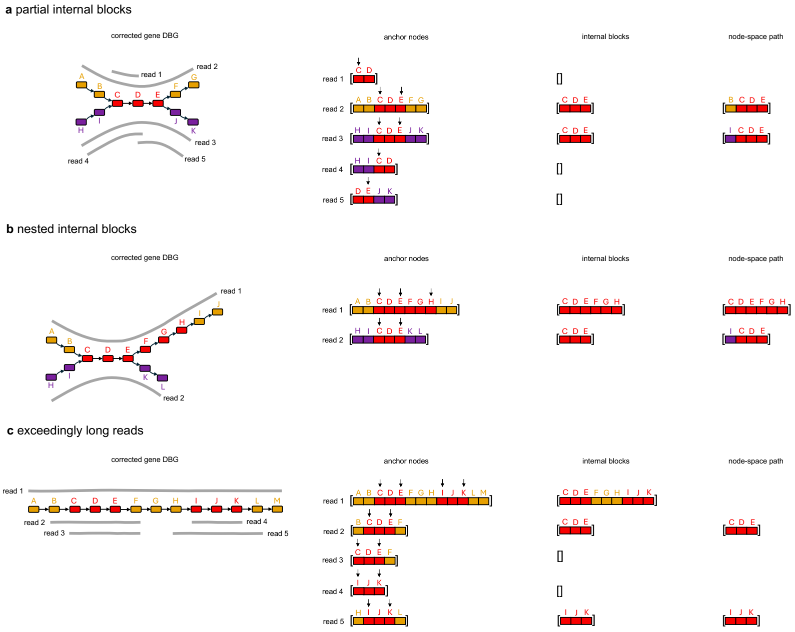


**Fig. S10** Examples of complex paths that can be expected in the read assignment stage of Amira and the resulting anchor nodes, internal blocks and final node-space paths.
